# Passive epidemiological surveillance in wildlife in Costa Rica identifies pathogens of zoonotic and conservation importance

**DOI:** 10.1101/2021.12.17.473220

**Authors:** Fernando Aguilar-Vargas, Tamara Solorzano-Scott, Mario Baldi, Elías Barquero-Calvo, Ana Jiménez-Rocha, Carlos Jiménez, Marta Piche-Ovares, Gaby Dolz, Bernal León, Eugenia Corrales-Aguilar, Mario Santoro, Alejandro Alfaro-Alarcón

**Author notes:** Contributed equally to this work: Fernando Aguilar-Vargas, Tamara Solorzano-Scott.

## Abstract

Epidemiological surveillance systems for pathogens in wild species have been proposed as a preventive measure for epidemic events. These systems can minimize the detrimental effects of an outbreak, but most importantly, passive surveillance systems are the best adapted to countries with limited resources. Therefore, the present study aims to evaluate the technical and infrastructural feasibility to establish this type of scheme in Costa Rica targeting thedetection of pathogens of zoonotic and conservation importance in wildlife. Between 2018 and 2020, 85 carcasses of free-ranging vertebrates were admitted for post mortem analysis and complementary laboratory analysis, representing a solid basis for the implementation of a passive surveillance system for wildlife diseases in the country. However, we encounter during this research significant constraints that affected the availability of carcasses for analysis, mainly related to the initial identification of cases, detection biases towards events in populated- or easily accessible-areas with nearby located wildlife management centers, further associated with financial disincentives, and limited local logistics capacity. Thus resulting in the exclusion of some geographic regions of the country. This epidemiological surveillance scheme allowed us to estimate the general state of health of the country’s wildlife, establishing the cause of death of the analyzed animals as follows: (i) 46 (54.1%) traumatic events, (ii) 23 (27.1%) infectious agents, (iii) two (2.4%) degenerative illness, (iv) three (3.5%) presumably poisoning, and (v) in 11 (12.9%)undetermined. It also allowed the detection of pathogens such as, canine distemper virus, *Klebsiella pneumoniae, Toxoplasma gondii, Trypanosoma* spp., *Angiostrongylus* spp., *Dirofilaria* spp., *Baylisascaris* spp., among others. As well as recognizing the circulation of these pathogens around national territory and also on those analyzed species. This strategy is crucial in geographical regions defined as critical for the appearance of diseases due to their great biodiversity and social conditions.

## Introduction

Zoonotic diseases represent one of the major burdens to society (both locally and globally), and a direct threat to public health, conservation, and human welfare programs [1,2]. Moreover, zoonotic diseases are responsible for causing significant economic losses, distorting social patterns, and negatively impacting human development indices, with a strong impact on developing countries [3–6]. A current example is the COVID-19 pandemic, that evidences the inadequacy of the infrastructures and diagnostic facilities/techniques to ensure surveillance, for early detection of potentially zoonotic agents in wildlife and for establishing control measures and mitigation [7].

Wildlife populations act as reservoirs for numerous pathogens, many of which can affect both human and animal health though with different degrees. Wildlife play various roles within the epidemiology of diseases participating in their spread and maintenance [8–10]. These roles assign wildlife the important function of sentinels of the ecosystems’ health, and allows the early detection of alterations in the environment; as well as, the distribution, the re-emergence or emergence of certain pathogens in a specific region [11,12].

Tropical countries (including Costa Rica) are among the areas of most extraordinary natural diversity with concomitant high diversity of pathogens and thus, a high potential for disease emergence [13,14]. The risk posed by wildlife as possible natural reservoirs, especially in geographic regions such as Costa Rica, a crossing point between North America and South America, is stressed by migratory movements that influence the appearance of diseases [15,16]. This risk has increased drastically because of anthropogenic pressures linked to over-exploitation of natural resources and increase of land use that in turn increases the possibility of contact between wildlife, domestic animals, and humans [17,18].

One of the preventive strategies against the risk of epidemic events, promoted by the World Organization for Animal Health (OIE) and World Health Organization (WHO) is to increase the efforts to establish early detection mechanisms for pathogens, of both zoonotic and conservation importance, via epidemiological monitoring systems, and emphasizing the need to develop robust surveillance and control programs for diseases in free-ranging species [7,19,20]. As early as 2012, the World Bank stated that preventive investments in the “ONE HEALTH” approach through veterinary and public health services could mean significant savings in response to the zoonotic outbreaks the world faces annually [21]. This statement reinforces the idea that investment in programs of epidemiological surveillance using wildlife as sentinels may be an alternative measure to prevent or predict epidemic events, which could minimize the economic impact of an outbreak.

One of the first steps to know the health status of the wildlife in a region is monitoring through passive surveillance, which identifies the causes of morbidity and mortality in a range of species based on their pathological profiles through post-mortem examinations. This approach offers advantages such as profitability and the ability to carry out a convenience sampling by taking advantage of the established infrastructure and obtaining a sustainable tool that allows understanding the emerging potential of different pathogens. Furthermore, when these schemes are set in the long term, it has been proven to provide the core information for decision-making and policies, regulations, and strategies establishment, prioritizing disease prevention, even when the sampling is biased and with incomplete geographic coverage [22–25].

Significant efforts have been made in Latin American to improve epidemiological surveillance systems aimed at animal health, however, there is still the need to optimize these schemes [26,27]. For example, according to the U.S. Department of Agriculture, Costa Rica has the infrastructure and maintains adequate surveillance programs to detect and control zoonotic diseases in farm animals [28]. However, it does not contemplate local wildlife within its scheme as it should be [29].

Several pathogens such as zoonotic parasites, vector-borne diseases, and direct transmission viruses have been identified in Costa Rican wildlife [30–41]. Nonetheless, without being contemplated in a routine systematic wildlife surveillance program, this information cannot reveal the local distribution of these infectious agents or the general health status of wildlife. And it only reflects the urgency of establishing a constant and sustainable surveillance diseases system, where aspects such as the pathogens zoonotic in wildlife are contemplated, and open the door to further research to know the eco-epidemiology of these agents at the local level.

Countries with limited resources (LMIC) such as Costa Rica, face severe financial and logistical restrictions to monitor the health and circulation of pathogens in wildlife. Nevertheless, the animal health system should, in the short term, establish at least a passive epidemiological surveillance in wildlife. Therefore, during our research a pilot scheme for surveillance of the general health status of wildlife was implemented, aiming to evaluate the technical and infrastructural feasibility to establish sustainably of this type of scheme in Costa Rica. Eighty-five carcasses of wild species were analyzed, detecting zoonotic pathogens, such us *Klebsiella pneumoniae, Toxoplasma gondii, Baylisascaris spp*. and pathogens of conservation importance in wildlife, such us canine distemper virus, among others. Furthermore, our research allowed to know the circulating pathogens in the analyzed species and their distribution in the national territory. This demonstrated that there is the logistical capacity to implement this surveillance program even when we experience some limitations.

## Material and Methods

### Statement of Ethics

All samples were obtained from dead wildlife (found dead in the field or euthanized after veterinary care in specialized centers). The study was approved by the Ministry of Environment and Energy (MINAE) (wildlife authority) through permit (R-SINAC-PNI-ACLAC-039), and with the support of the animal health authority, the National Animal Health Service through the office (SENASA-DG-0277-18).

### Notification and receptionof cases

Between 2018 and 2020, officials from the wildlife management centers, veterinarians from the National Animal Health Service (SENASA), and wildlife officials from the National Wildlife Service (SINAC) reported cases, and voluntarily sent specimens from different localities in the country to the Wildlife Program of SENASA or directly to Pathology Department of the Escuela de Medicina Veterinaria, Universidad Nacional. The carcasses used in this study were all from free-ranging vertebrates with any apparent clinical condition or after death due to any associated disease or trauma. Basic information was requested for every sample submission: geographic location, the standard and scientific name of the animal, clinical signs, and any information considered relevant to the case, following the scheme recommended by the OIE for the notification of cases for disease surveillance system in wild animals [20,42]. All carcasses were shipped fresh under refrigerated conditions or stored at -20 °C for a maximum period of one week before shipping.

### Pathological analysis

The carcasses received were classified by their autolysis degree according to an established scale of one to five [43]. Thus, ranging from a fresh carcass or recently dead animal (grade 1) to advanced decomposition (grade 4) and partial, mummified carcasses or skeletal remains (grade 5). Only carcasses with grades 1 to 3 were included in the study for post-mortem analysis and tissue sampling [44]. Therefore, 96 specimens were received, of which, 85 were admitted in the study. These were divided by taxonomic class into birds and mammals. The mammal class was subdivided into taxonomic groups. The geographical distribution and the taxa admitted are shown in S1 Fig.

All morphological findings were recorded. In addition, tissue samples were taken for routine histopathological and microbiological analysis as required. Tissue samples for histopathology were processed based on standard routing protocols [44].

### Detection of different infectious agents

#### Virus Detection

Molecular methods were used for the detection of different viral agents. All molecular methods were done in the presence of a positive and a negative control. The samples analyzed were fresh tissues collected in a sterile manner during post-mortem analysis. In addition, we performed RNA extraction using the commercial kit DNeasy Blood and Tissue (QIAGEN, Venlo, The Netherlands), following the manufacturer’s recommendations. The methods used and the samples collected are specified in Table 1.

**Table 1.**
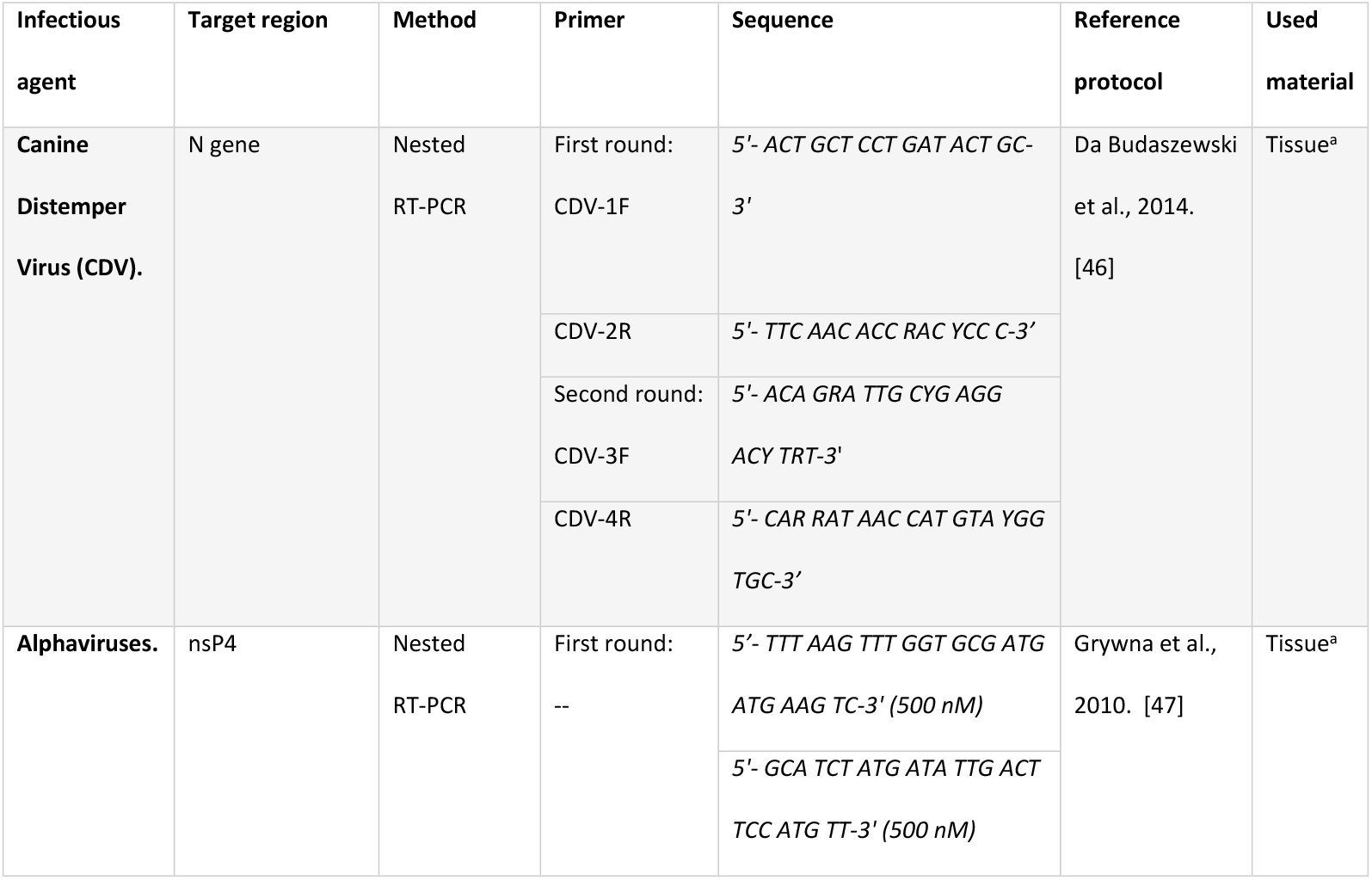
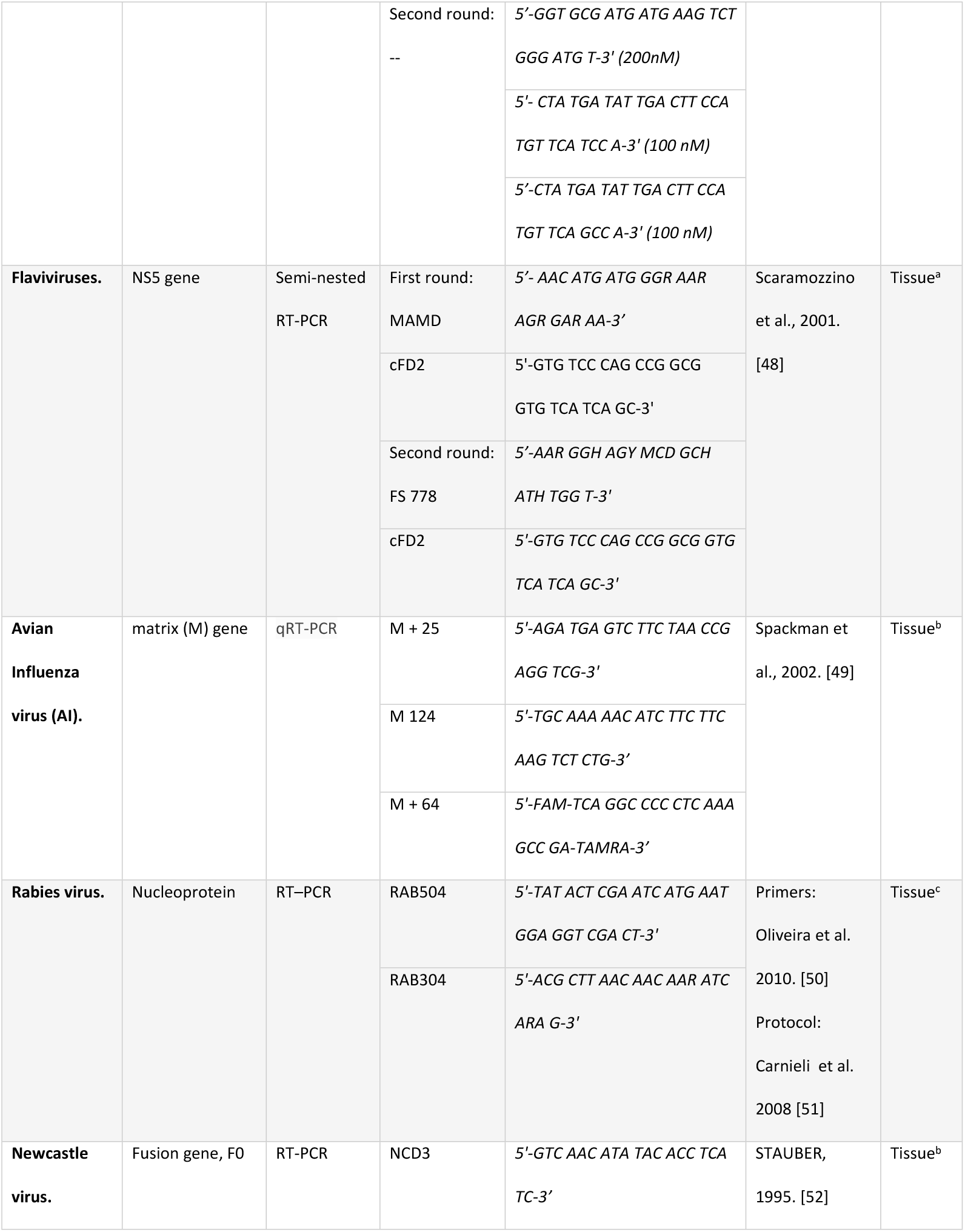
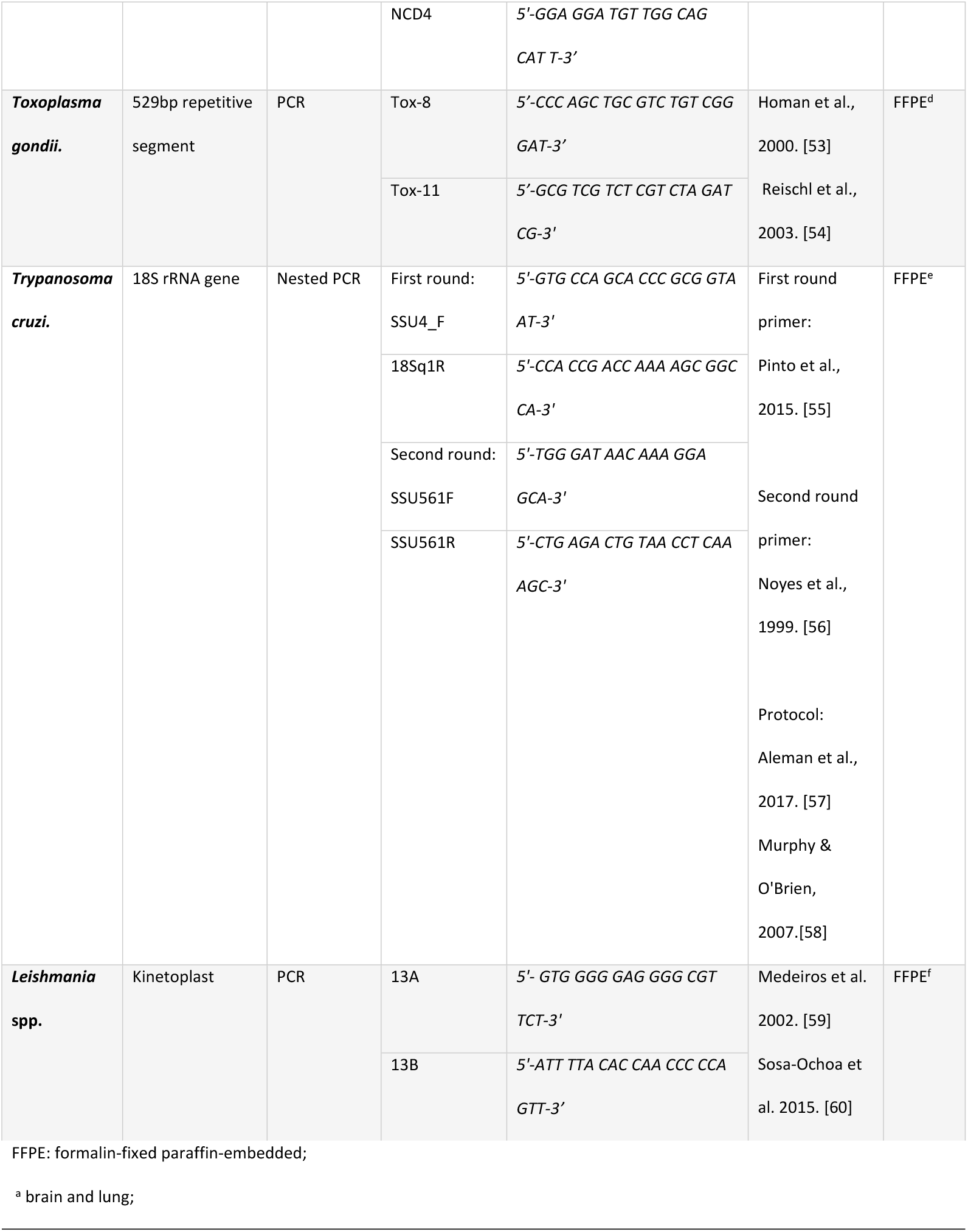
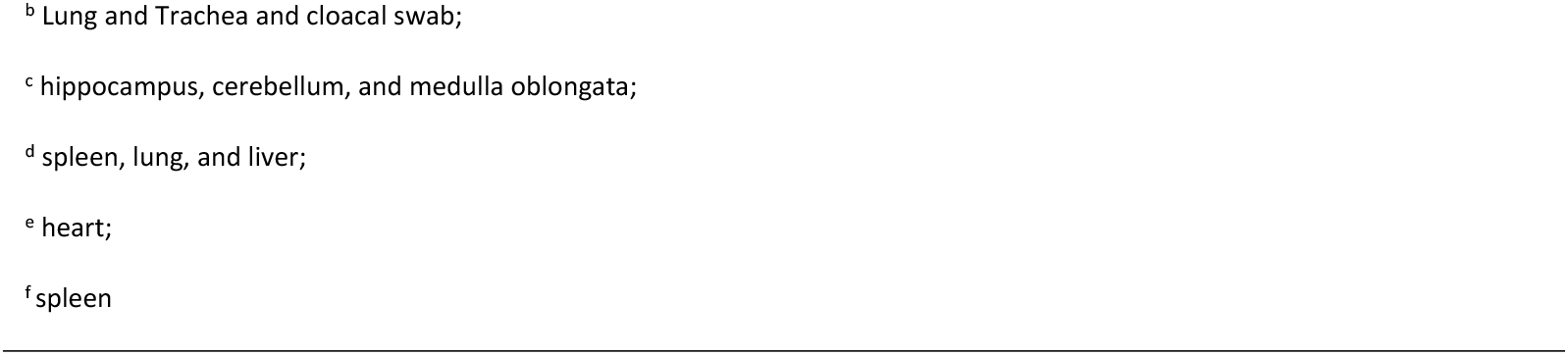
Molecular techniques for the detection of viral agents and protozoa.

#### Protozoa Detection

Confirmation was performed using molecular techniques for pathogen identification when a previous presumptive protozoa presence was established in the histological study. All molecular methods were done in the presence of a positive and a negative control. Tissue samples previously embedded in paraffin were used for this purpose. The deparaffinization procedure was done using xylol washes following the method recommended to perform DNA extraction from the tissue [45]. According to the manufacturer’s instructions, we performed DNA extraction using the commercial kit DNeasy Blood and Tissue (QIAGEN, Venlo, The Netherlands). The methods used and the samples collected are specified in Table 1.

#### Bacteriological detection

Tissue samples from animals with inflammatory processes (suppurative or abscesses) were cultured following standard bacteriological procedures. For bacterial isolation, samples were inoculated on non-selective and selective agar media. Significant bacterial growth was identified using the automated VITEK-2 Compact system, software version 8.02 (bioMérieux, Marcy l’Etoile, France). VITEK test cards for Gram-negative [GN], Gram-positive [GP], and anaerobes [ANC] were used for identification according to the manufacturer’s instructions.

#### Parasites identification

All parasites recovered from animals of the two investigated classes were washed with physiological saline, preserved in alcohol, acetic acid, and formalin (AFA) solution, and subjected to identification to genus level [61]. Physical and morphometric characteristics were recognized after fixation and clarification with Hoyer’s solution [62–64]. In addition, processed cestodes were stained with dilute Harris’ hematoxylin solution.

### Geocoding and spatial analysis

Each case was geocoded using the latitude and longitude generated by GPS of the point where the specimen was found by field personnel. When the GPS was not available, they were geocoded using the latitude and longitude of the approximate location where they were found, and this was generated by Google Earth Pro v7.3 (2021, Google Inc.). The georeferenced points of each sample admitted created a map using ArcGIS 10.7 (ERSI) (S1 Fig).

## Results

### Distribution of cases for age, sex, taxonomic classification and sender

Of the 85 specimens admitted to the study, there was an age distribution of 23 (27.1%) young animals and 62 (72.9%) adults. The sex distribution was 48 (56.5%) males and 37 (43.5%) females. According to the taxonomic class, we received nine (10.5%) specimens of the birds and 76 (89.5%) of the mammals. The notification of cases was made by SINAC and the Animal Health Service SENASA, with 21 (24.7%) of the cases and 64 (75.3%) by wildlife rescue centers. The biological data of the admitted specimens are shown in S2 Table. Furthermore, cases of two mass fatality events were received. In the first case, field officials reported the death of hundreds of pelicans, of which we received two specimens and they were found positive for flaviviruses. The second case was the death of several carnivores who tested positive for the canine distemper virus.

### Distribution of causes of death and injuries according to etiology

The distribution of the presumptive cause of death according pathological findings corresponded to 46 (54.1%) associated with traumatic events (mainly roadkill and electrocution), 23 (27.1%) with fatalities directly related to infectious agents, and 2 (2.4%) with degenerative disease. Additionally, in 3 (3.5%) cases, the death was presumptively associated with intoxication, and in 11 (12.9%), the cause of death was not determined. Of the individuals with a cause of traumatic death, 31 (67.4%) concomitantly presented lesions associated with an infectious etiology (24 with parasites, three with bacteria, one with protozoa and four with multiple microorganisms). Of the specimens with an infectious cause of death, ten presented lesions associated with viruses, five with parasites, two with protozoa, one with bacteria, and five presented lesions associated with multiple etiologies.

Of the identified lesions by analyzed organs (gross and microscopic injuries), 199 (74.8%) were associated with infectious processes, and 67 (25.2%) were associated with non-infectious processes (46 traumatic, five toxic, four degenerative diseases, and twelve non-relevant focal lesions). According to the predominant inflammatory infiltrate of the infectious processes, presumptively 69 (34.7%) lesions were associated with protozoa and parasites (histiocytic and eosinophilic infiltrate, respectively), 55 (27.6%) were linked with viral infections (lymphoplasmacytic infiltrate), and 36 (18.1%) were related to bacterial processes (suppurative infiltrate). Examples of these inflammatory lesions is observed in Fig 1 (see legend). In 39 lesions, the infectious etiology was not investigated because they were considered non-relevant focal lesions. The distribution of lesions identified by taxon and etiology are described in Table 2. The description of the pathological findings identified by species is detailed in S2 Table.

**Fig 1.**
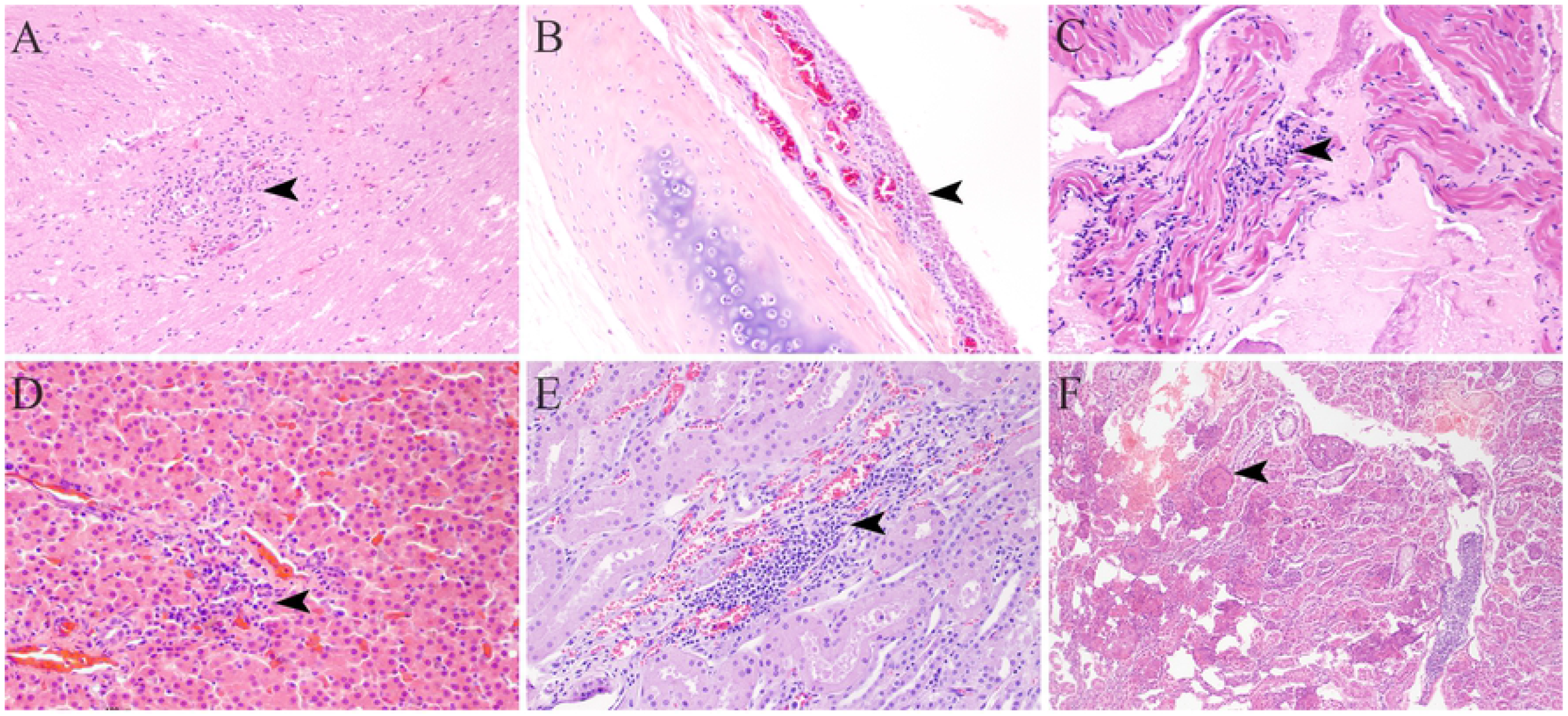
Frequent pathological findings associated with inflammatory processes in wild animals. A) Brain (*Procyon lotor*-raccoon). Perivascular lymphoplasmacytic encephalitis associated with CDV infection (Arrowhead; H&E 200x). B) Trachea (*Pelecanus occidentalis*-brown pelican). Ulcerative haemorrhagic pyogranulomatous tracheitis with pseudomembrane formation (arrowhead: H&E 200x). C) Heart (*Didelphis marsupialis*-opossum). Lymphoplasmacytic myocarditis associated with *Trypanosoma cruzi* amastigotes (Arrowhead; H&E 200x). D) Liver (*Pelecanus occidentalis-*brown pelican). Lymphoplasmacytic and necrotizing hepatitis associated with flavivirus infection (arrowhead; H&E 200x) E) Kidney (*Canis latrans*-coyote). Lymphoplasmacytic interstitial nephritis (arrowhead; H&E 200x). F) Kidney (*Jabiru mycteria*-jabiru). Necrotizing granulomatous glomerulonephritis with crystal formation (arrowhead; H&E 100x).

**Table 2.**
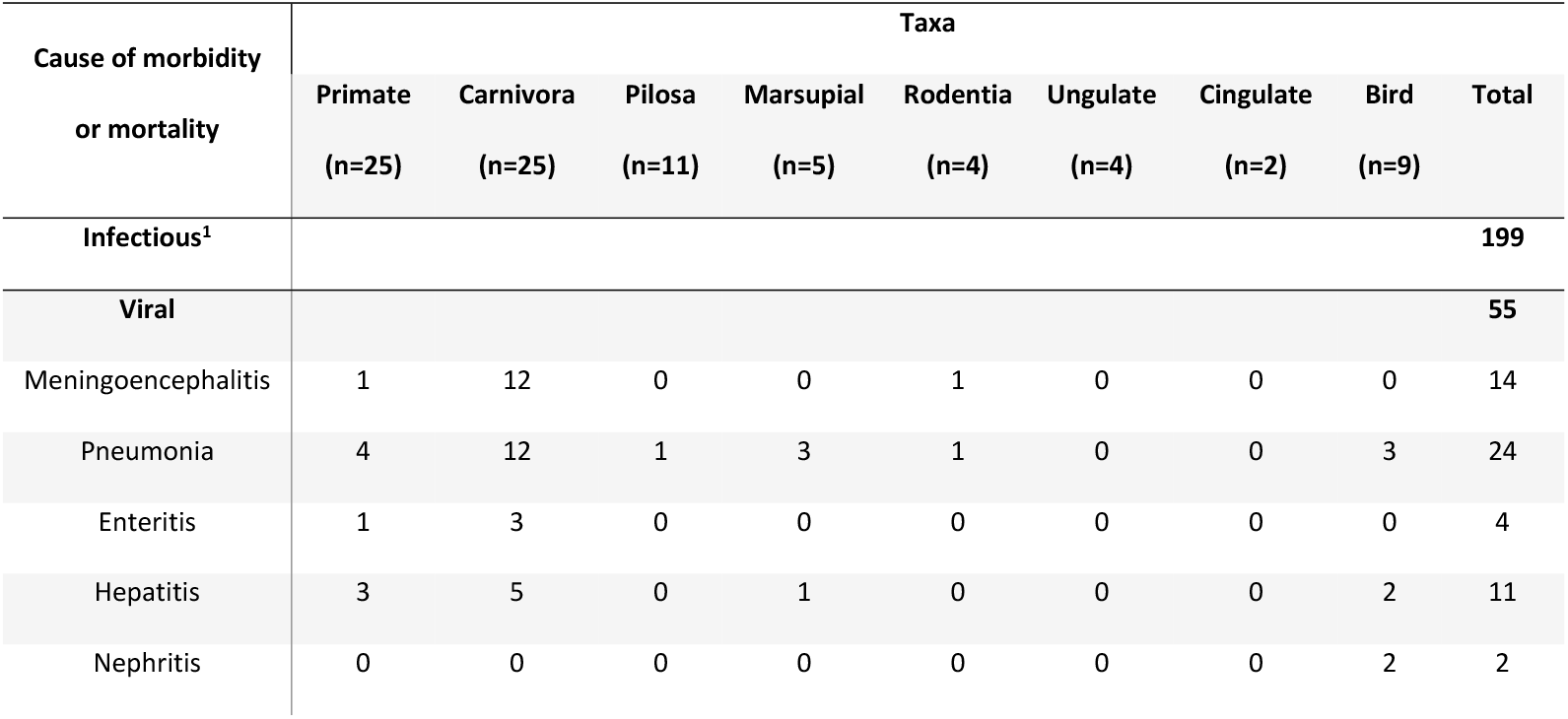
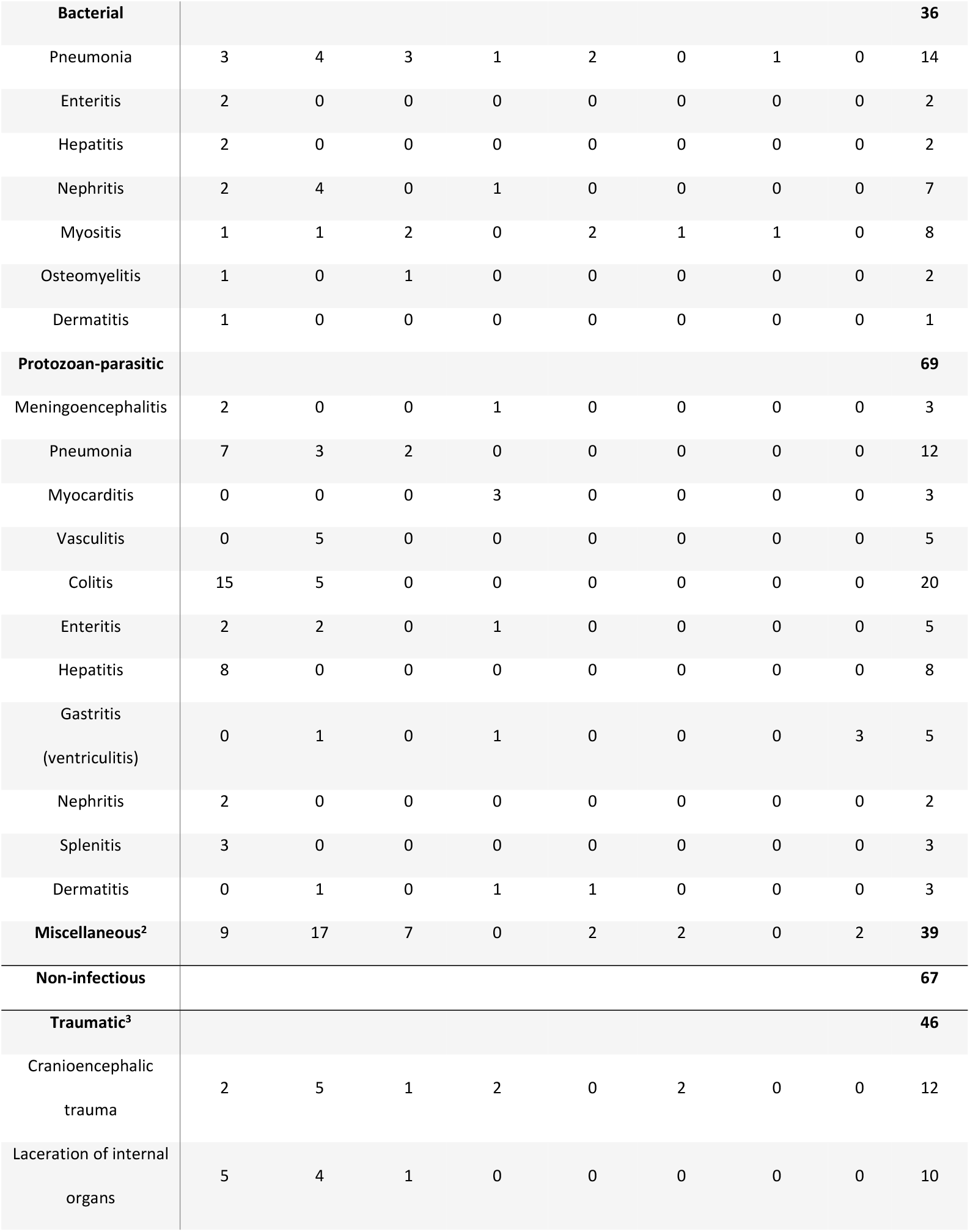
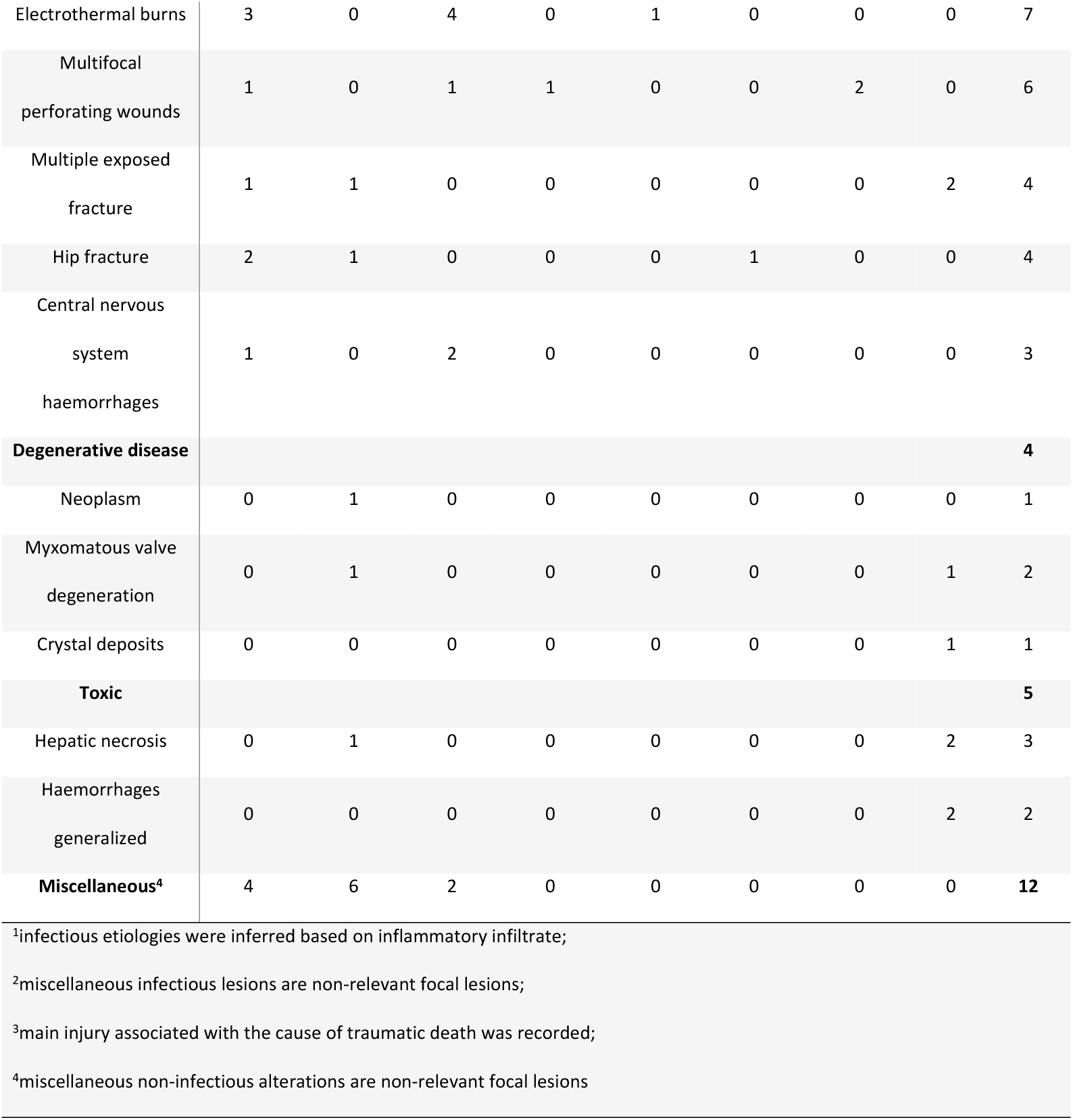
Main morphological diagnoses identified in free-ranging wild animals analyzed in the years 2018-2020.

Of the lesions associated with infectious etiology, in 44 (22.1%), the causal agent was not possible to be determined. In 21 of these lesions, a predominantly lymphoplasmacytic inflammatory infiltrate was observed, suggestive of viral etiology (11 pneumonia, five hepatitis, four encephalitis). In 18 lesions, the inflammatory infiltrate was suppurative, suggestive of bacterial etiology (five pneumonia, four nephritis, four myositis, one dermatitis). Five lesions had a predominantly histiocytic infiltrate suggestive of the presence of protozoa (n=2 hepatitis, n=2 colitis, n=1 splenitis). All parasites associated with lesions were identified in the complementary analysis.

### Identification of viral, protozoan, bacterial and parasitic infectious agents by taxonomic group

Ten virus, seven protozoa, and seven bacteria were identified in mammalian specimens. In 22 cases, these pathogens were involved with lesions or systemic disease, of which, 19 were directly associated with the cause of death of mammals. Only *Sarcocystis* spp. detected in two cases was an incidental finding. Additionally, 38 mammals had internal parasites (23 different genera). Multi-parasitosis was observed in 13 (17.1%) of the cases. Parasites such as *Prosthenorchis* spp. (n=15), *Angiostrongylus* spp. (n=6), and *Cilycospirura* spp. (n=1) were responsible for severe parasitosis with systemic disease. Some of the lesions such as pyogranulomatous and eosinophilic meningoencephalitis, pyogranulomatous abscessing bronchopneumonia and eosinophilic gastritis associated with infectious agents are observed in Fig 2 (see legend). In 69% (n=43) of the mammalian cases, infectious agents with zoonotic potential such as *Klebsiella pneumoniae, Toxoplasma gondi*i, *Angiostrongylus* spp. were identified. The etiological agents identified by taxonomic groups and the number of specimens analyzed are specified in Table 3.

**Fig 2.**
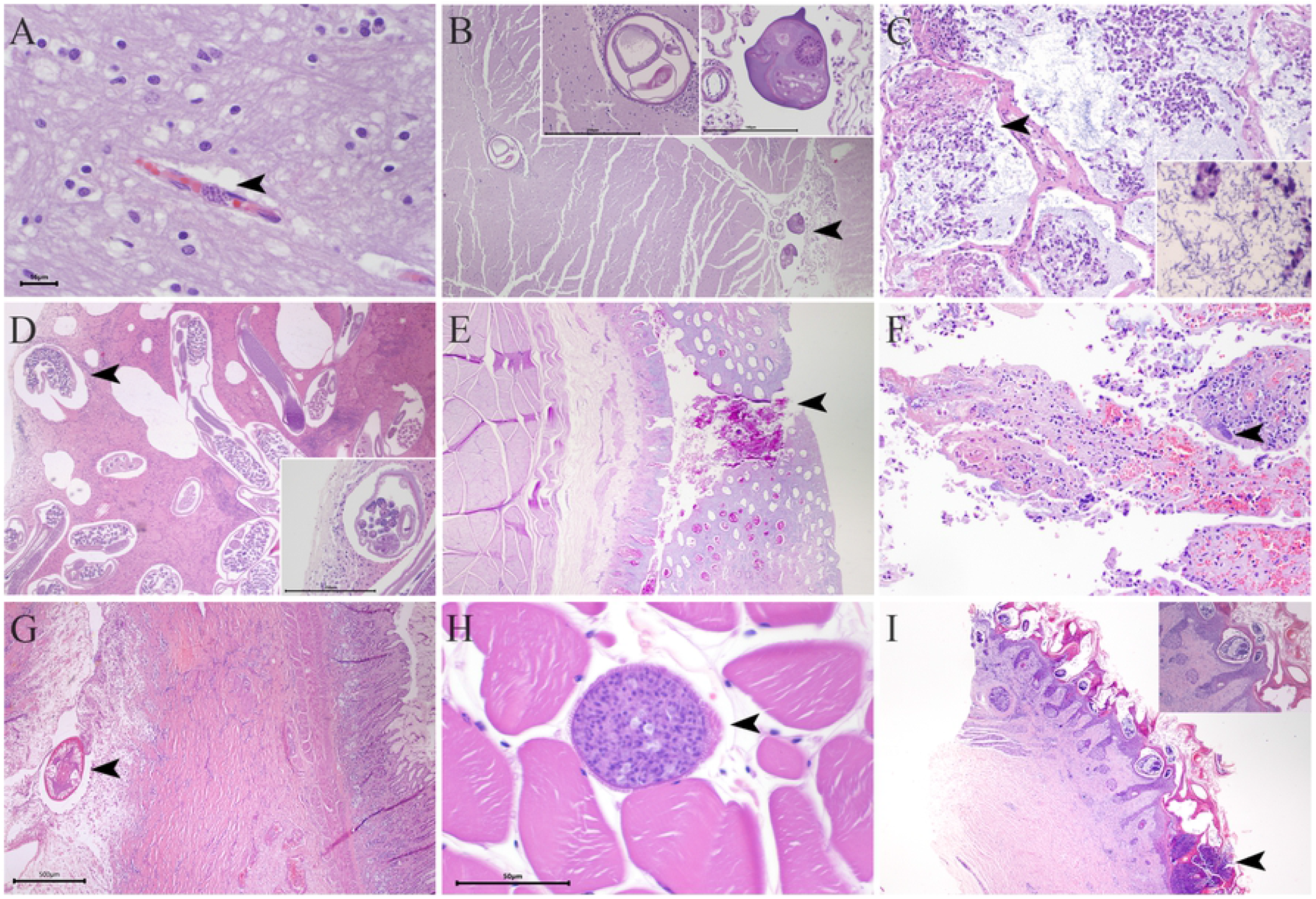
Infectious agents in lesions identified in wild animals. A) Brain (*Alouatta palliata*-howler monkey). Presence of protozoan pseudocysts in the blood vessel, morphology compatible with *Toxoplasma gondii* (arrowhead; H&E 600x). B) Brain (*Didelphis marsupialis*-opossum). Pyogranulomatous and eosinophilic meningoencephalitis associated with *Angiostrongylus* spp. (arrowhead; H&E 400x). Inset: Nematode magnification (H&E 200x & 400x). C) Lung (*Alouatta palliata*-howler monkey). Pyogranulomatous abscessing bronchopneumonia associated with *Klebsiella pneumoniae* (arrowhead; H&E 200x). Inset: Bacterial aggregates magnification (H&E 600x). D) Lung (*Cebus imitator*-white-faced monkey). Pyogranulomatous and eosinophilic pleuro-bronchopneumonia associated to multiple *Filariopsis* spp. nematodes (arrowhead; H&E 40x). Inset: Nematode magnification (H&E 200x). E) Ventricle (*Pelecanus occidentalis-* brown pelican). Pyogranulomatous ulcerative and eosinophilic ventriculitis associated with multiple *Contracaecum* spp. nematodes (arrowhead; H&E 40x). F) Jejunum (*Ateles geoffroyi*-spider monkey). Pyogranulomatous and necrotizing jejunitis associated with *Staphylococcus aureus* infection (arrowhead; H&E x200). G) Stomach (*Herpailurus yagouaroundi*-jaguarundi). Eosinophilic gastritis associated with multiple *Cylicospirura* spp. nematodes (arrowhead; H&E 40x). H) Skeletal muscle (*Nasua narica*-coati). Presence of protozoan cyst, morphology consistent with *Sarcocystis* spp (arrowhead; H&E 200x). I) Skin (*Sphiggurus mexicanus*-porcupine) Pyogranulomatous and eosinophilic dermatitis associated with massive infestation of *Sarcoptex* spp. (arrowhead; H&E 400x). Inset: Mites magnification (H&E 100x).

**Table 3:** Number of infectious agents tested and positive in mammals according to etiology.

| Etiological agent |  | Mammalian taxonomic groups (positives) |  |  |  |  |  |  |
| --- | --- | --- | --- | --- | --- | --- | --- | --- |
|  |  | Primate | Carnivora | Pilosa | Marsupial | Rodentia | Ungulate | Cingulate |
| **Viral** | Canine Distemper Virus (n=18) | 0 | 10 | 0 | 0 | 0 | 0 | 0 |
|  | Alphaviruses (n=9) | 0 | 0 | 0 | 0 | 0 | 0 | 0 |
|  | Flaviviruses (n=9) | 0 | 0 | 0 | 0 | 0 | 0 | 0 |
|  | Influenza virus (n=8) | 0 | 0 | 0 | 0 | 0 | 0 | 0 |
|  | Rabies virus (n=76) | 0 | 0 | 0 | 0 | 0 | 0 | 0 |
| **Bacterial** | *Clostridium perfringens* (n= 18) | 0 | 0 | 0 | 0 | 0 | 1 | 0 |
|  | *Escherichia coli* (n=18) | 1 | 0 | 0 | 0 | 0 | 0 | 0 |
|  | *Klebsiella pneumoniae* (n=18) | 1 | 0 | 0 | 0 | 0 | 0 | 0 |
|  | *Trueperella pyogenes*. (n=18) | 0 | 0 | 0 | 0 | 1 | 0 | 0 |
|  | *Staphylococcus aureus* (n=18) | 1 | 1 | 1 | 0 | 0 | 0 | 0 |
|  | *Mycobacterium* spp. (n=18) | 0 | 0 | 0 | 0 | 0 | 0 | 0 |
| **Protozoan** | *Toxoplasma gondii* (n=4) | 2 | 0 | 0 | 0 | 0 | 0 | 0 |
|  | *Trypanosoma* spp. (n=14) | 0 | 0 | 0 | 3 | 0 | 0 | 0 |
|  | *Leishmania* spp. (n=8) | 0 | 0 | 0 | 0 | 0 | 0 | 0 |
|  | *Sarcocystis* spp. (n=5) | 0 | 1 | 0 | 0 | 0 | 1 | 0 |
| **Parasitic<sup>1</sup>** | *Angiostrongylus* spp. | 0 | 5 | 0 | 1 | 0 | 0 | 0 |
|  | *Dirofilaria* spp. | 0 | 4 | 0 | 0 | 0 | 0 | 0 |
|  | *Dipetalonema* spp. | 5 | 0 | 2 | 0 | 0 | 0 | 0 |
|  | *Gnathostoma* spp. | 0 | 0 | 0 | 1 | 0 | 0 | 0 |
|  | *Baylisascaris* spp. | 0 | 1 | 0 | 0 | 0 | 0 | 0 |
|  | *Ancylostoma* spp. | 0 | 1 | 0 | 0 | 0 | 0 | 0 |
|  | *Cylicospirura* spp. | 0 | 1 | 0 | 0 | 0 | 0 | 0 |
|  | *Prosthenorchis* spp. | 10 | 5 | 0 | 0 | 0 | 0 | 0 |
|  | *Macracanthorhynchus* spp. | 0 | 1 | 0 | 0 | 0 | 0 | 0 |
|  | *Spirometra* spp. | 0 | 2 | 0 | 0 | 0 | 0 | 0 |

n= Number tested;
<sup>1</sup> Only zoonotic parasites are shown

All birds submitted were evaluated for the presence of the (n=9) virus, two of the birds, which were involved in an episode of high mortality during the study period, were positive for flaviviruses. Additionally, three birds had internal parasites (2 different genera). Most of the pathogens identified were directly associated as the cause of the death of birds and linked with lesions illustrated in Fig 2. Only *Procyrnea* spp. identified in one case was an incidental finding. In 22.2% (n=2) of the birds cases, infectious agents with zoonotic potential such as *Contracaecum* spp. were identified. The etiological agents identified in birds and the number of samples analyzed are specified in Table 4.

**Table 4:** Number of infectious agents tested and positive in birds according to etiology.

| Etiological agent |  | Birds |
| --- | --- | --- |
| Viral diseases |  | Positive |
|  | Alphaviruses (n=3) | 0 |
|  | Flaviviruses (n=3) | 2 |
|  | Influenza virus (n=9) | 0 |
|  | Newcastle virus (n=9) | 0 |
| Bacterial diseases | *Clostridium perfringens* (n=1) | 0 |
|  | *Escherichia coli* (n=1) | 0 |
|  | *Klebsiella pneumoniae* (n=1) | 0 |
|  | *Salmonella* spp. (n=1) | 0 |
|  | *Staphylococcus aureus* (n=1) | 0 |
| **Parasites diseases** | *Contracaecum* spp. | 2 |

n= number of tested;
<sup>1</sup> only zoonotic parasites are shown

### Geospatial distribution of detected infectious agents and their accumulation by geographic region

We established a distribution of the most frequently identified infectious agents in the analyzed specimens (Fig 3). First, a wide distribution of zoonotic parasites was evidenced in the country. Then, there was an accumulation in the Central Pacific region of specimens with acanthocephaliasis (12 with *Prosthenorchis* spp., one with *Macracanthorhynchus* spp.), and an accumulation of specimens with gastrointestinal nematodes in the great metropolitan area and tourist areas of Guanacaste (six with *Angiostrongylus* spp., one with *Baylisascaris* spp., one with *Ancylostoma* spp.). Additionally, vector-borne diseases occurred exclusively in specimens from coastal regions and altitudes less than 300 meters above sea level (11 with filariae, two with flaviviruses). The CDV present in carnivores from various areas of the country did not show a specific pattern of distribution (n=10). It is important to note that the Caribbean-south and southern part of the country do not present any pattern because there was no shipment of samples from this area, a caveat of this scheme of passive surveillance. The analyzed specimens associated with these infectious agents can be observed in S1 Fig.

**Fig 3.**
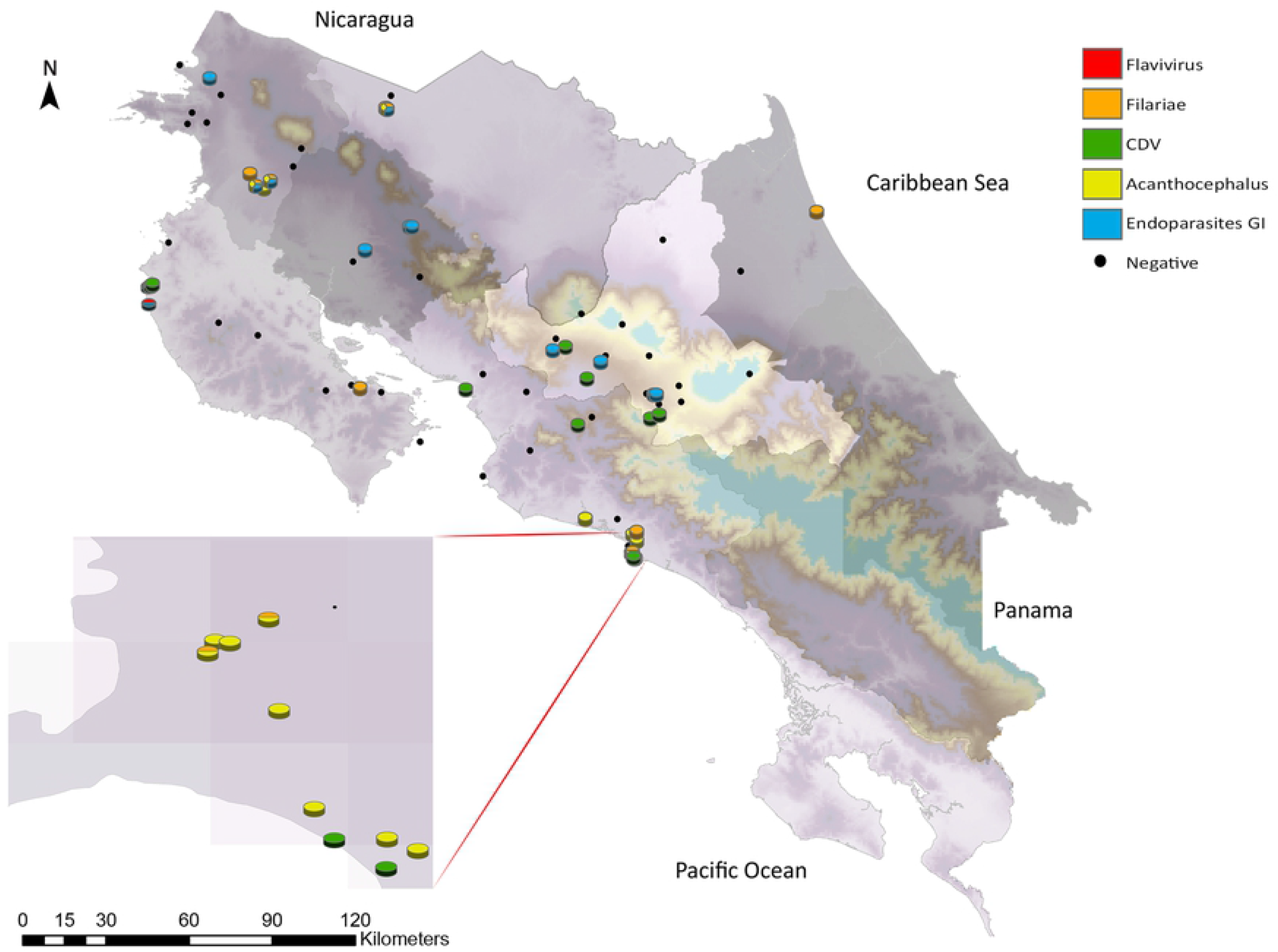
Geographical distribution of the most frequently identified infectious agents in the referred specimens. The individuals reported as negative were depicted even though the infectious agent was not detected in the complementary analyzes or no lesions suggestive of the disease were found in the pathological analysis.

## Discussion

The schemes aimed to develop epidemiological surveillance of infectious agents in free-living vertebrates have proven to be a fundamental tool in monitoring pathogens of zoonotic importance [65–67]. These surveillance systems are even more critical in geographical areas where high rates of biodiversity are prominent [7]. For example, Costa Rica is economically dependent on its ecotourism services, and its fauna is one of its most important assets [68]. However, currently, there is no epidemiological surveillance system directed to wildlife and to study outbreaks or any other health event involving them.

Evaluating the viability of implementing a passive surveillance system should be essential for a country considered a “hotspot” for the appearance or emergence of new infectious agents associated with its biodiversity and fauna characteristics [14,18]. However, some authors have established obstacles to implementing this type of system. These obstacles are mainly related to bureaucratic restrictions, financial disincentives, lack of legislation for data collection, and willingness to cooperate between agencies [69,70]. We encounter during this research those same obstacles for implementing surveillance systems in wildlife animals in Costa Rica.

Centers specialized in wildlife disease surveillance and health have shown that it is possible to establish robust and sustainable disease surveillance systems with broad coverage and diagnostic capacity [25,65,66,71]. However, factors such as logistics, rescue centers location, and dependence on voluntary staff excluded the participation of some geographic regions of the country (remote or difficult-to-access areas), affecting the efficiency and sustainability of surveillance system implemented in our research. These findings are consistent with other studies, where significant constraints hindered the availability of carcasses for analysis [72,73].

Reporting and dispatching of carcasses by national wildlife and animal health authorities were lower than rescue centers. These patterns agree with previous reports indicating that notification of wildlife mortality or morbidity generally depends on the initial detection of cases by the general public [22,74–76]. As a result, cases are biased towards events in populated or easily accessible areas with nearby wildlife management centers, this added to the logistic capacity (storage, packaging, and shipping) of some of these centers, would explain the higher dispatch of cases. An example of these biases is the accumulation of reported cases in the Central Pacific region.

Wildlife management centers often report data with similar mortalities between birds and mammals [67,76]. However, in our study, there is a higher number of mammalians received. This difference is probably related to the fact that the surveillance systems in those countries actively include avian influenza surveillance, leading to a higher reporting of birds than our study. In our case, there is no surveillance system for avian influenza in wild birds by the local veterinary authority, and the suspected cases of this virus are confined to the local poultry production systems [71]. On the other hand, most of the birds found were no longer suitable for the study, thus underestimating the number of birds that die for various reasons [77].

Regarding the mammals, two taxa presented a higher representation (carnivores and primates). These data can be associated with the fact that they are medium to large-sized animals and with a more significant contact of these species with human environments, facilitating the recognition of morbidities and mortalities [23]. Human proximity with these mammals enabled the detection of infectious diseases, which is highly represented in these groups [78,79].

The distribution of the causes of death established in our research is consistent with most studies, highlighting traumatic events (anthropogenic effect) as the leading cause of death in these wild species [73,80–82]. Most of the traumas correspond to vehicular collisions, with a high percentage reported in urban areas or roads that cross or pass near forest areas. However, the number of deceased animals may be underestimated since not all cases are reported or recorded, much less referred to the laboratory since the condition of the corpses is not suitable for the study [83,84].

In addition, most of the traumatic cases presented some pre-existing infectious pathology. Free-living animals are frequently exposed to infectious agents naturally, so it is common in post-mortem analysis to find incidental lesions associated [82,85,86]. In wild animals highly exposed to human conditions, factors such as climate change and anthropogenic impact could become stressful stimuli, which facilitate these infectious agents to evolve into severe disease and increase the risk of suffering a traumatic event [82,87]. However, it is difficult to establish a clear association [88,89].

Some of these identified infectious agents cause foodborne illness in human populations. Examples of these are *Clostridium* spp., *Toxoplasma* sp. and *Sarcocystis* spp. Although the law prohibits hunting in Costa Rica, there is the illegal consumption of wild animal meat. Consequently, this type of practice might favor the transmission of infectious agents and lead to local outbreaks or maintain circulating virulent strains in local human populations [90–93].

Our study shows the presence of potentially zoonotic bacterial infectious agents classified as emerging diseases in some regions [94–96]. The most relevant are *Klebsiella pneumoniae, Escherichia coli*, and *Staphylococcus aureus*, which were associated with primary disease in some of the analyzed specimens. The direct or indirect contact occurs through the handling of these wild animals, thus facilitating the transmission, which evidences a latent risk. In addition, these bacteria currently top the list of infectious agents with antibiotic resistance genes, making them considered within antimicrobial surveillance schemes [97,98]. Furthermore, cases of antimicrobial resistance have already been demonstrated in Costa Rica with other bacteria in wild animals in urban environments, thus reflecting the need for a wildlife pathogen surveillance scheme to consider active antimicrobial resistance monitoring [31,99].

Zoonotic vector-borne diseases arise when there is a conjunction of spatial-temporal factors and other variables (reservoirs, climate, susceptible population, among others) [100]. Environmental conditions in tropical regions favor these diseases that significantly impact public health and are recognized as agents with epidemic potential in Latin America [101–103]. We identified many primates and carnivores with infectious agents of vector transmission, for example, *Dirofilaria* spp. and *Dipetalonema* spp. mainly present in low-lying areas (CoastLine). These regions are already defined as endemic areas for these parasites in domestic animals [104,105]. Nevertheless, detecting this type of agent in a jungle cycle reveals a potential risk to public health in places with a high rate of visiting tourists in Costa Rica. This risk is reinforced by reports of the health system, which showed at least three disease cases in humans associated with *Dirofilaria immitis* and isolated cases of subcutaneous filariasis [106–108].

Similarly, several Latin American countries are considered endemic to various diseases caused by arboviruses [109]. In Costa Rica, other studies have identified the stationary circulation of this type of agent in humans and animals in various regions [33,110,111]. The cases of arbovirus-flaviviruses detected in our research are consistent with the high circulation of this type of virus in Costa Rica [33,111]. Furthermore, this agent was associated with a mass mortality of pelicans during the conduct of our research. Detecting virus-related mortalities such as West Nile in wild birds (as it was possibly our case) allows early alerts. It has been shown that there is a higher risk of exposure for human populations close to the regions where mortalities of wild birds occur [112,113].

The canine Distemper virus (CDV) (genus: morbillivirus) is a pernicious infectious agent with a global distribution that affects at least 20 families of mammals. Especially susceptible are carnivores of all species [114,115]. Endemic CDV outbreaks have been reported anecdotally throughout Costa Rica and America in dog populations and, more recently, sporadic outbreaks in wild carnivores of urban and suburban areas in the country have been recorded [116,117]. CDV was identified in our studies, reflecting the relevance of this virus in the role of spillover towards carnivore species and possibly the implications of a spillback towards susceptible or non-vaccinated domestic canines [117–119]. Costa Rica has a domestic dog population of ∼250 thousand animals, mainly located in urban areas. Herd immunity data in this population is uncertain, especially for dogs without an owner or in non-urban areas, where owners neglect these vaccines. Indeed, this poses a risk to wild carnivores, especially in urban areas with susceptible canine populations. Furthermore, the possibility of transmission of this virus to other species beyond carnivores is a hypothesis that has been investigated [120]. Given the high diversity of vertebrates present in Costa Rica, this virus should be considered within epidemiological surveillance programs.

This study did not detect rabies virus infections. This findings are supported by previous studies in wild animals in Costa Rica [32]. However, human and productive animal fatalities have been reported associated with rabies infections, which stresses the relevance of its continuous monitoring [38,121]. A similar situation applies to Newcastle and Influenza virus. In our samples, none of the birds showed evidence of disease or associated clinical signs. However, due to the sanitary status (declared itself free) of Costa Rica for these avian viral agents and the risk for national poultry production, it is advisable to establish monitoring in any event of mortality of wild birds in the country [122,123].

The gastrointestinal and pulmonary parasites detected in this study are relevant for public health and wildlife conservation programs. For instance, the nematodes *Baylisascaris* spp., *Ancylostoma* spp., and *Cylicospirura* spp. were detected in mammalian species located in densely populated areas. Furthermore, we identified parasites transmitted by water or food of aquatic origin (such as the cestode *Spirometra* spp. and the nematodes *Contracaecum* spp. and *Gnathostoma* spp.) mainly in rural areas of the country’s northern region. In this region, fishing and rivers for recreational, irrigation, and consumption purposes are common, showing possible contamination in both ways [124,125].

The last two reports of human angiostrongyliasis in Costa Rica have shown 12.9% seropositivity in the screening test, the majority were children under the age of ten who reside in San José, which is the province with the highest number of human samples [126,127]. In this study, six specimens of mammals infected by *Angiostrongylus* spp. were identified, mainly in the *Nasua narica* species, from recreational parks in cities in the country’s northern region. This species has been established as the definitive host in the parasite’s life cycle [128]. The high degree of positive cases suggests a high prevalence of the parasite in the mammalian reservoir. This result evidences the urgency of expanding sampling with better diagnostic techniques in children and wild carnivores from these regions. The same situation happens with the number of cases detected with acanthocephalans (*Prosthenorchis* spp., *Macracanthorhynchus* spp.) in the Central Pacific region. There is no information on the real prevalence in animal populations, nor are there samplings that allow detecting cases in humans in this region, despite the zoonotic risk previously mentioned [129,130].

Finally, we could not identify the causative agent of lesions in some of the samples analyzed. However, histological changes suggest the presence of an infectious agent. Although we analyzed samples for the main circulating infectious agents in Costa Rica, no conclusive data was obtained for some of them. Ranges of 17-22% have been reported in pathological studies in wild species, where the causative agent of the disease cannot be determined, mainly associated with the degree of autolysis and the diagnostic complexity [25,73,82,85]. These results are consistent with the percentages of an absence of identification of the etiological agent in our samples. These results show that further work is necessary to develop robust diagnostic techniques for wild animals and efforts and incentives financed by government authorities in the surveillance of pathogens in wildlife through the consistent implementation of new generation metagenomics [131–134].

Most of the pathogens detected in our study have already been previously identified in wild animals in Costa Rica, and the detection of an infectious agent in a wild specimen does not necessarily imply disease or affect wild populations [30,33,41,128]. However, monitoring the general health status of wild animals over time allows us to know the circulation and behavior of these pathogens, as well as to provide an early warning of epidemic events. This information can be used by health authorities together with a preventive strategy and a ONE HEALTH approach to address zoonotic diseases, facilitating more specific public health interventions, implementing measures to reduce the risk of spread [25,66,73].

This study was performed as a pilot and was the first structured attempt to test the establishment of a passive epidemiological surveillance scheme for diseases in wild vertebrates. However, it highlights the necessity of an inter-institutional and trans-institutional commitment with the sustainability over time of this surveillance scheme focused on the benefits beyond the economic part. For example, this work allowed us to estimate the general health status of the country’s wildlife and know the distribution of pathogens in the national territory. This information is critical in regions established as hotspots for the emergence of infectious diseases due to their great biodiversity and social conditions [18].

## Acknowledgments

The authors express deep appreciation to the National Animal Health Service (SENASA) and National Wildlife Service (SINAC) for their support and cooperation provided in this study and the wildlife management centers for the notification of specimens, especially DVM. Isabel Hagnauer Barrantes from Zooave rescue center, DVM. Martha Cordero Salas from Las Pumas rescue center and DVM. Carmen Soto Valverde from the Kids Saving the Rainforest rescue center, for their particular interest and participation. Also, the collaboration provided by the technicians of the laboratories of the Escuela de Medicina Veterinaria of the Universidad Nacional is appreciated.

## Author Contributions

**Conceptualization: A.A**.

**Data curation: A.A**., **F.A**., **T.S**., **M.B**.

**Formal analysis: A.A**., **F.A**., **T.S**., **M.B**.

**Funding Acquisition: A.A**.

**Investigation: A.A**., **F.A**., **T.S**.

**Project administration: A.A**.

**Resources: A.A**., **T.S**., **E.B**., **A.J**., **C.J**., **M.P**., **G.D**., **M.S**.

**Supervision: A.A**.

**Software: A.A**., **F.A**., **T.S**., **M.B**.

**Visualization: A.A**., **F.A, T.S**.

**Writing – original draft: A.A**., **F.A**., **T.S**., **M.B**., **E.B**.

**Writing – review & editing: A.A.A**., **F.A**., **T.S**., **M.B**., **E.B**., **A.J**., **C.J**., **M.P**., **G.D**., **B.L**., **E.C.A**., **M.S**.

**All authors have read and agreed to the published version of the manuscript**.

**Conflicts of Interest: the authors declare no conflict of interest**.

## Supporting information

**S1 Fig. Geographical distribution of the analyzed specimens and number of individuals by taxon**.

**S2 Table. Biological data and more representative pathological findings of the analyzed specimens**.

## Notes

### Competing Interest Statement

The authors have declared no competing interest.

## References

1. Morens DM, Folkers GK, Fauci AS. The challenge of emerging and re-emerging infectious diseases. Nature. 2004; 430:242–9. doi: 10.1038/nature02759 PMID: 15241422.

2. Bloom DE, Cadarette D. Infectious Disease Threats in the Twenty-First Century: Strengthening the Global Response. Front Immunol. 2019; 10:549. doi: 10.3389/fimmu.2019.00549 PMID: 30984169.

3. CEPAL. América Latina y el Caribe ante la pandemia del COVID-19 Efectos económicos y sociales. Chile: ONU 2020. Available from: https://repositorio.cepal.org/handle/11362/45337.

4. PNUD. Evaluación económica inicial de los efectos de covid-19 y el alcance de las opciones de política en Costa Rica. Síntesis para la discusión y análisis de políticas. Costa Rica: 19 programa de las naciones unidas para el desarrollo 2020. Available from: https://www.cr.undp.org/content/costarica/es/home/library/evaluacion-economica-inicial-de-los-efectos-de-covid-19-y-alcanc.html.

5. Shepard DS, Coudeville L, Halasa YA, Zambrano B, Dayan GH. Economic impact of dengue illness in the Americas. Am J Trop Med Hyg. 2011; 84:200–7. doi: 10.4269/ajtmh.2011.10-0503 PMID: 21292885.

6. Zohrabian A, Meltzer MI, Ratard R, Billah K, Molinari NA, Roy K, et al. West Nile virus economic impact, Louisiana, 2002. Emerging Infect Dis. 2004; 10:1736–44. doi: 10.3201/eid1010.030925 PMID: 15504258.

7. Petrovan SO, Aldridge DC, Bartlett H, Bladon AJ, Booth H, Broad S, et al. Post COVID-19: a solution scan of options for preventing future zoonotic epidemics. Biol Rev Camb Philos Soc. 2021. Epub 2021/07/07. doi: 10.1111/brv.12774 PMID: 34231315.

8. Viana M, Mancy R, Biek R, Cleaveland S, Cross PC, Lloyd-Smith JO, et al. Assembling evidence for identifying reservoirs of infection. Trends Ecol Evol (Amst). 2014; 29:270–9. doi: 10.1016/j.tree.2014.03.002 PMID: 24726345.

9. Hassell JM, Begon M, Ward MJ, Fèvre EM. Urbanization and Disease Emergence: Dynamics at the Wildlife-Livestock-Human Interface. Trends Ecol Evol (Amst). 2017; 32:55–67. doi: 10.1016/j.tree.2016.09.012 PMID: 28029378.

10. Haydon DT, Cleaveland S, Taylor LH, Laurenson MK. Identifying reservoirs of infection: a conceptual and practical challenge. Emerging Infect Dis. 2002; 8:1468–73. doi: 10.3201/eid0812.010317 PMID: 12498665.

11. Reif JS. Animal sentinels for environmental and public health. Public Health Rep. 2011; 126 Suppl 1:50–7. doi: 10.1177/00333549111260S108 PMID: 21563712.

12. Kowalewski MM, Salzer JS, Deutsch JC, Raño M, Kuhlenschmidt MS, Gillespie TR. Black and gold howler monkeys (Alouatta caraya) as sentinels of ecosystem health: patterns of zoonotic protozoa infection relative to degree of human-primate contact. Am J Primatol. 2011; 73:75–83. doi: 10.1002/ajp.20803 PMID: 20084672.

13. Jones KE, Patel NG, Levy MA, Storeygard A, Balk D, Gittleman JL, et al. Global trends in emerging infectious diseases. Nature. 2008; 451:990–3. doi: 10.1038/nature06536 PMID: 18288193.

14. Walsh MG, Sawleshwarkar S, Hossain S, Mor SM. Whence the next pandemic? The intersecting global geography of the animal-human interface, poor health systems and air transit centrality reveals conduits for high-impact spillover. One Health. 2020; 11:100177. Epub 2020/10/08. doi: 10.1016/j.onehlt.2020.100177 PMID: 33052311.

15. Engering A, Hogerwerf L, Slingenbergh J. Pathogen-host-environment interplay and disease emergence. Emerg Microbes Infect. 2013; 2:e5. Epub 2013/06/02. doi: 10.1038/emi.2013.5 PMID: 26038452.

16. Altizer S, Bartel R, Han BA. Animal migration and infectious disease risk. Science. 2011; 331:296–302. doi: 10.1126/science.1194694 PMID: 21252339.

17. White RJ, Razgour O. Emerging zoonotic diseases originating in mammals: a systematic review of effects of anthropogenic land-use change. Mamm Rev. 2020. Epub 2020/06/02. doi: 10.1111/mam.12201 PMID: 32836691.

18. Allen T, Murray KA, Zambrana-Torrelio C, Morse SS, Rondinini C, Di Marco M, et al. Global hotspots and correlates of emerging zoonotic diseases. Nat Commun. 2017; 8:1124. Epub 2017/10/24. doi: 10.1038/s41467-017-00923-8 PMID: 29066781.

19. World Health Organization. Anticipating Emerging Infectious Disease Epidemics. Switzerland: WHO 2015. Available from: https://apps.who.int/iris/bitstream/handle/10665/252646/WHO-OHE-PED-2016.2-eng.pdf.

20. World Organisation for Animal Health. Training manual on surveillance and international reporting of diseases in wild animals second cycle. Workshop for OIE National Focal Points for Wildlife. Paris: OIE 2015. Available from: https://www.oie.int/fileadmin/Home/eng/Internationa_Standard_Setting/docs/pdf/WGWildlife/A_Training_Manual_Wildlife_2.pdf.

21. International Bank for Reconstruction and Development. People, pathogens and our planet. The Economics of One Health. Washington DC: The World Bank 2012. Available from: https://openknowledge.worldbank.org/bitstream/handle/10986/11892/691450ESW0whit0D0ESW120PPPvol120web.pdf?sequence=1&isAllowed=y.

22. Sleeman JM. Strategies for Wildlife Disease Surveillance. United States: U.S. Geological Survey National Wildlife Health Center 2012. Available from: https://digitalcommons.unl.edu/cgi/viewcontent.cgi?article=1981&context=usgsstaffpub.

23. Guberti V. Surveillance, monitoring and surveys of wildlife diseases: a public health and conservation approach. Italy: Hystrix, the Italian Journal of Mammalogy 2014. Available from: http://www.italian-journal-of-mammalogy.it/Surveillance-monitoring-and-surveys-of-wildlife-diseases-a-public-health-and-conservation,77207,0,2.html.

24. Lamarque F. Le reseau SAGIR, reseau national de suivi sanitaire de la faune sauvage française. Paris: Epidémiol. et santé anim 2000. Available from: https://www.researchgate.net/profile/Marc_Artois/publication/237744255_Le_reseau_SAGIR_reseau_national_de_suivi_sanitaire_de_la_faune_sauvage_francaise/links/00b7d529709d2110a0000000/Le-reseau-SAGIR-reseau-national-de-suivi-sanitaire-de-la-faune-sauvage-francaise.pdf.

25. Leighton FA, Wobeser GA, Barker IK, Daoust PY, Martineau D. The Canadian Cooperative Wildlife Health Centre and surveillance of wild animal diseases in Canada. Can Vet J. 1997; 38:279–84. Available from: https://www.ncbi.nlm.nih.gov/pmc/articles/PMC1576906/pdf/canvetj00090-0025.pdf.

26. OPS. Veterinary public health: progress report on the secretariat’s compliance with the mandates of the paho governing bodies. México: Organización Panamericana de la Salud 2005. Available from: https://iris.paho.org/bitstream/handle/10665.2/40542/rimsa14-03-e.pdf?sequence=1&isAllowed=y.

27. Rojas H, Romero JR. Where to next with animal health in Latin America? The transition from endemic to disease-free status. Rev - Off Int Epizoot. 2017; 36:331–48. doi: 10.20506/rst.36.1.2633 PMID: 28926004.

28. USDA. Report on the Review of Costa Rica’s Animal Health Statuses. Estados Unidos: United States Department of Agriculture 2019. Available from: https://www.aphis.usda.gov/import_export/downloads/costarica-status-review.pdf.

29. World Organisation for Animal Health. Informe de Misión Piloto Evaluación PVS “Una Salud”. Paris: OIE 2011. Available from: https://www.oie.int/fileadmin/Home/eng/Support_to_OIE_Members/pdf/InterimReport-Costa_Rica.pdf.

30. Baldi M, Alvarado G, Smith S, Santoro M, Bolaños N, Jiménez C, et al. Baylisascaris procyonis Parasites in Raccoons, Costa Rica, 2014. Emerging Infect Dis. 2016; 22:1502–3. doi: 10.3201/eid2208.151627 PMID: 27433741.

31. Baldi M, Barquero Calvo E, Hutter SE, Walzer C. Salmonellosis detection and evidence of antibiotic resistance in an urban raccoon population in a highly populated area, Costa Rica. Zoonoses Public Health. 2019; 66:852–60. Epub 2019/07/29. doi: 10.1111/zph.12635 PMID: 31359623.

32. Baldi M, Hernández-Mora G, Jimenez C, Hutter SE, Alfaro A, Walzer C. Leptospira Seroprevalence Detection and Rabies Virus Absence in an Urban Raccoon (Procyon lotor) Population in a Highly Populated Area, Costa Rica. Vector Borne Zoonotic Dis. 2019; 19:889–95. Epub 2019/08/13. doi: 10.1089/vbz.2019.2444 PMID: 31407956.

33. Chaves A, Piche-Ovares M, Ibarra-Cerdeña CN, Corrales-Aguilar E, Suzán G, Moreira-Soto A, et al. Serosurvey of Nonhuman Primates in Costa Rica at the Human-Wildlife Interface Reveals High Exposure to Flaviviruses. Insects. 2021; 12. Epub 2021/06/15. doi: 10.3390/insects12060554 PMID: 34203687.

34. Dolz G, Chaves A, Gutiérrez-Espeleta GA, Ortiz-Malavasi E, Bernal-Valle S, Herrero MV. Detection of antibodies against flavivirus over time in wild non-human primates from the lowlands of Costa Rica. PLoS One. 2019; 14:e0219271. doi: 10.1371/journal.pone.0219271 PMID: 31276532.

35. Dubey JP, Morales JA, Sundar N, Velmurugan GV, González-Barrientos CR, Hernández-Mora G, et al. Isolation and genetic characterization of Toxoplasma gondii from striped dolphin (Stenella coeruleoalba) from Costa Rica. J Parasitol. 2007; 93:710–1. doi: 10.1645/GE-1120R.1 PMID: 17626370.

36. Fuentes-Ramírez A, Jiménez-Soto M, Castro R, Romero-Zuñiga JJ, Dolz G. Molecular Detection of Plasmodium malariae/Plasmodium brasilianum in Non-Human Primates in Captivity in Costa Rica. PLoS One. 2017; 12:e0170704. doi: 10.1371/journal.pone.0170704 PMID: 28125696.

37. González K, Calzada JE, Saldaña A, Rigg CA, Alvarado G, Rodríguez-Herrera B, et al. Survey of wild mammal hosts of cutaneous leishmaniasis parasites in panamá and costa rica. Trop Med Health. 2015; 43:75–8. Epub 2014/12/06. doi: 10.2149/tmh.2014-30 PMID: 25859156.

38. Hutter SE, Brugger K, Sancho Vargas VH, González R, Aguilar O, León B, et al. Rabies in Costa Rica: Documentation of the Surveillance Program and the Endemic Situation from 1985 to 2014. Vector Borne Zoonotic Dis. 2016; 16:334–41. Epub 2016/03/16. doi: 10.1089/vbz.2015.1906 PMID: 26982168.

39. Patiño W LC, Monge O, Suzán G, Gutiérrez-Espeleta G, Chaves A. Molecular Detection of Mycobacterium avium avium and Mycobacterium genavense in Feces of Free-living Scarlet Macaws (Ara macao) in Costa Rica. J Wildl Dis. 2018; 54:357–61. doi: 10.7589/2017-05-124 PMID: 29286261.

40. Pérez-Gómez G, Jiménez-Rocha AE, Bermúdez-Rojas T. Parásitos gastrointestinales de aves silvestres en un ecosistema ribereño urbano tropical en Heredia, Costa Rica. RBT. 2018; 66:788. doi: 10.15517/rbt.v66i2.33409.

41. Rojas-Jiménez J, Morales-Acuña JA, Argüello-Sáenz M, Acevedo-González SE, Yabsley MJ, Urbina-Villalobos A. Histopathological findings of infections caused by canine distemper virus, Trypanosoma cruzi, and other parasites in two free-ranging White-nosed Coatis Nasua narica (Carnivora: Procyonidae) from Costa Rica. J Threat Taxa. 2021; 13:17521–8. doi: 10.11609/jott.5907.13.1.17521-17528.

42. World Organisation for Animal Health. Training manual on wildlife diseases and surveillance. Workshop for OIE National Focal Points for Wildlife. Paris: OIE 2010. Available from: https://www.oie.int/fileadmin/Home/eng/Internationa_Standard_Setting/docs/pdf/WGWildlife/A_Training_Manual_Wildlife.pdf.

43. McAloose D, Colegrove KM, Newton AL. Wildlife Necropsy. Pathology of Wildlife and Zoo Animals. Elsevier; 2018. pp. 1–20.

44. Woodford MH, Bengis RG, Keet DF. Post-mortem procedures for wildlife veterinarians and field biologists. Paris: OIE; 2000.

45. Körbler T, Gršković M, Dominis M, Antica M. A simple method for RNA isolation from formalin-fixed and paraffin-embedded lymphatic tissues. Experimental and Molecular Pathology. 2003; 74:336–40. doi: 10.1016/s0014-4800(03)00024-8.

46. Da Budaszewski RF, Pinto LD, Weber MN, Caldart ET, Alves CDBT, Martella V, et al. Genotyping of canine distemper virus strains circulating in Brazil from 2008 to 2012. Virus Res. 2014; 180:76–83. Epub 2013/12/24. doi: 10.1016/j.virusres.2013.12.024 PMID: 24370870.

47. Grywna K, Kupfer B, Panning M, Drexler JF, Emmerich P, Drosten C, et al. Detection of all species of the genus Alphavirus by reverse transcription-PCR with diagnostic sensitivity. J Clin Microbiol. 2010; 48:3386–7. doi: 10.1128/JCM.00317-10 PMID: 20504990.

48. Scaramozzino N, Crance JM, Jouan A, DeBriel DA, Stoll F, Garin D. Comparison of flavivirus universal primer pairs and development of a rapid, highly sensitive heminested reverse transcription-PCR assay for detection of flaviviruses targeted to a conserved region of the NS5 gene sequences. J Clin Microbiol. 2001; 39:1922–7. doi: 10.1128/JCM.39.5.1922–1927.2001 PMID: 11326014.

49. Spackman E, Senne DA, Myers TJ, Bulaga LL, Garber LP, Perdue ML, et al. Development of a real-time reverse transcriptase PCR assay for type A influenza virus and the avian H5 and H7 hemagglutinin subtypes. J Clin Microbiol. 2002; 40:3256–60. doi: 10.1128/JCM.40.9.3256-3260.2002 PMID: 12202562.

50. Oliveira RdN, Souza SP de, Lobo RSV, Castilho JG, Macedo CI, Carnieli P, et al. Rabies virus in insectivorous bats: implications of the diversity of the nucleoprotein and glycoprotein genes for molecular epidemiology. Virology. 2010; 405:352–60. doi: 10.1016/j.virol.2010.05.030 PMID: 20609456.

51. Carnieli P, Fahl WdO, Castilho JG, Oliveira RdN, Macedo CI, Durymanova E, et al. Characterization of Rabies virus isolated from canids and identification of the main wild canid host in Northeastern Brazil. Virus Res. 2008; 131:33–46. Epub 2007/09/21. doi: 10.1016/j.virusres.2007.08.007 PMID: 17889396.

52. Stauber. N. Detection of Newcastle disease virus in poultry vaccines using the polymerase chain reaction and direct sequencing of amplified cDNA. Vaccine. 1995; 13:360–4. doi: 10.1016/0264-410X(95)98257-B.

53. Homan W, Vercammen M, Braekeleer J de, Verschueren H. Identification of a 200-to 300-fold repetitive 529 bp DNA fragment in Toxoplasma gondii, and its use for diagnostic and quantitative PCR1Note: Nucleotide sequence data reported in this paper have been submitted to GenBankTM database with the accession number AF146527 (Toxoplasma gondii genomic repetitive 529 bp fragment).1. International Journal for Parasitology. 2000; 30:69–75. doi: 10.1016/s0020-7519(99)00170-8.

54. Reischl U, Bretagne S, Krüger D, Ernault P, Costa J-M. Comparison of two DNA targets for the diagnosis of Toxoplasmosis by real-time PCR using fluorescence resonance energy transfer hybridization probes. BMC Infect Dis. 2003; 3:7. doi: 10.1186/1471-2334-3-7 PMID: 12729464.

55. Pinto CM, Ocaña-Mayorga S, Tapia EE, Lobos SE, Zurita AP, Aguirre-Villacís F, et al. Bats, Trypanosomes, and Triatomines in Ecuador: New Insights into the Diversity, Transmission, and Origins of Trypanosoma cruzi and Chagas Disease. PLoS One. 2015; 10:e0139999. Epub 2015/10/14. doi: 10.1371/journal.pone.0139999 PMID: 26465748.

56. Noyes H, Stevens J, Teixeira M, Phelan J, Holz P. A nested PCR for the ssrRNA gene detects Trypanosoma binneyi in the platypus and Trypanosoma sp. in wombats and kangaroos in Australia1. International Journal for Parasitology. 1999; 29:331–9. doi: 10.1016/S0020-7519(98)00167-2.

57. Aleman A, Guerra T, Maikis TJ, Milholland MT, Castro-Arellano I, Forstner MRJ, et al. The Prevalence of Trypanosoma cruzi, the Causal Agent of Chagas Disease, in Texas Rodent Populations. Ecohealth. 2017; 14:130–43. Epub 2017/01/13. doi: 10.1007/s10393-017-1205-5 PMID: 28091763.

58. Murphy WJ, O’Brien SJ. Designing and optimizing comparative anchor primers for comparative gene mapping and phylogenetic inference. Nat Protoc. 2007; 2:3022–30. doi: 10.1038/nprot.2007.429 PMID: 18007639.

59. Medeiros ACR, Rodrigues SS, Roselino AMF. Comparison of the specificity of PCR and the histopathological detection of leishmania for the diagnosis of American cutaneous leishmaniasis. Braz J Med Biol Res. 2002; 35:421–4. doi: 10.1590/S0100-879X2002000400002 PMID: 11960189.

60. Sosa-Ochoa W, Morales Cortedano X, Argüello S, Zuniga C, Henríquez J, Mejía R, et al. Ecoepidemiología de la Leishmaniasis cutánea no ulcerada en Honduras. Ciencia y Tecnología. 2015:115–28. doi: 10.5377/rct.v0i14.1799.

61. Mehlhorn H. Methods to Diagnose Parasites. In: Mehlhorn H, editor. Animal Parasites. Cham: Springer International Publishing; 2016. pp. 23–32.

62. Mehlhorn H. Worms (Helminths). In: Mehlhorn H, editor. Animal Parasites. Cham: Springer International Publishing; 2016. pp. 251–498.

63. Mehlhorn H. Worms of Humans. In: Mehlhorn H, editor. Human Parasites. Cham: Springer International Publishing; 2016. pp. 135–298.

64. Salfelder K, Liscano TR de, Sauerteig E. Helminthic diseases. In: Salfelder K, Liscano TR de, Sauerteig E, editors. Atlas of Parasitic Pathology. Dordrecht: Springer Netherlands; 1992. pp. 96–172.

65. Childs JE, Krebs JW, Real LA, Gordon ER. Animal-based national surveillance for zoonotic disease: quality, limitations, and implications of a model system for monitoring rabies. Prev Vet Med. 2007; 78:246–61. Epub 2006/11/28. doi: 10.1016/j.prevetmed.2006.10.014 PMID: 17129622.

66. Moede Rogall G, Sleeman JM. The USGS National Wildlife Health Center: Advancing wildlife and ecosystem health. Reston, VA; 2017.

67. Craig Stephen. CWHC Annual report 2019/2020. Canada: Canadia Wildlife Health Cooperative 2020. Available from: http://www.cwhc-rcsf.ca/docs/annual_reports/2019_2020_CWHC_Annual_Report_EN.pdf?v=20201207.

68. Benavides Vindas S. El aporte del turismo a la economía costarricense: más de una década después. Econom y Socied. 2020; 25:1–29. doi: 10.15359/eys.25-57.1.

69. Stitt T, Mountifield J, Stephen C. Opportunities and obstacles to collecting wildlife disease data for public health purposes: results of a pilot study on Vancouver Island, British Columbia. Can Vet J. 2007; 48:83–7, 89-90. doi: 10.4141/cjas68-011 PMID: 17310627.

70. Don Bamunusinghage Nihal P, Dangolla A, Hettiarachchi R, Abeynayake P, Stephen C. Challenges and opportunities for wildlife disease surveillance in sri lanka. J Wildl Dis. 2020; 56:538–46. Epub 2020/01/09. doi: 10.7589/2019-07-181 PMID: 31917632.

71. Terrier M. Health monitoring of wildlife in france: SAGIR network and epidemiological monitoring of chiroptera rabies. Frence: Académie Vétérinaire de France 2006. Available from: https://core.ac.uk/download/pdf/15524538.pdf.

72. Forster G. Evaluating the feasibility of using data and samples provided by wildlife organisations to develop a national wildlife surveillance scheme in Costa Rica. Master Thesis, Univerity of London. 2020.

73. Pewsner M, Origgi FC, Frey J, Ryser-Degiorgis M-P. Assessing Fifty Years of General Health Surveillance of Roe Deer in Switzerland: A Retrospective Analysis of Necropsy Reports. PLoS One. 2017; 12:e0170338. Epub 1/19/2017. doi: 10.1371/journal.pone.0170338 PMID: 28103325.

74. Stallknecht DE. Impediments to wildlife disease surveillance, research, and diagnostics. Curr Top Microbiol Immunol. 2007; 315:445–61. doi: 10.1007/978-3-540-70962-6_17 PMID: 17848074.

75. Gourlay P, Decors A, Moinet M, Lambert O, Lawson B, Beaudeau F, et al. The potential capacity of French wildlife rescue centres for wild bird disease surveillance. Eur J Wildl Res. 2014; 60:865–73. doi: 10.1007/s10344-014-0853-9.

76. Cox-Witton K, Reiss A, Woods R, Grillo V, Baker RT, Blyde DJ, et al. Emerging infectious diseases in free-ranging wildlife-Australian zoo based wildlife hospitals contribute to national surveillance. PLoS One. 2014; 9:e95127. Epub 2014/05/01. doi: 10.1371/journal.pone.0095127 PMID: 24787430.

77. Balseiro A, Espí A, Márquez I, Pérez V, Ferreras MC, Marín JFG, et al. Pathological features in marine birds affected by the prestige’s oil spill in the north of Spain. J Wildl Dis. 2005; 41:371–8. doi: 10.7589/0090-3558-41.2.371 PMID: 16107672.

78. Han BA, Kramer AM, Drake JM. Global Patterns of Zoonotic Disease in Mammals. Trends Parasitol. 2016; 32:565–77. Epub 2016/06/14. doi: 10.1016/j.pt.2016.04.007 PMID: 27316904.

79. Plourde BT, Burgess TL, Eskew EA, Roth TM, Stephenson N, Foley JE. Are disease reservoirs special? Taxonomic and life history characteristics. PLoS One. 2017; 12:e0180716. Epub 2017/07/13. doi: 10.1371/journal.pone.0180716 PMID: 28704402.

80. Garcês A. A review of the mortality of wild fauna in Europe in the last century: the consequences of human activity. Journal of Wildlife and Biodiversity. 2020; 4:34–55. Available from: http://www.wildlife-biodiversity.com/article_37220_2f56660d6a06c893fdf3c6aced8de4e3.pdf.

81. Ascensão F, Desbiez ALJ, Medici EP, Bager A. Spatial patterns of road mortality of medium–large mammals in Mato Grosso do Sul, Brazil. Wildl Res. 2017; 44:135. doi: 10.1071/WR16108.

82. Navas-Suárez P, Díaz-Delgado J, Matushima ER, Fávero CM, am Sánchez Sarmiento, Sacristán C, et al. A retrospective pathology study of two Neotropical deer species (1995-2015), Brazil: Marsh deer (Blastocerus dichotomus) and brown brocket deer (Mazama gouazoubira). PLoS One. 2018; 13:e0198670. Epub 2018/06/07. doi: 10.1371/journal.pone.0198670 PMID: 29879222.

83. Carvajal V. Registro de mamíferos silvestres atropellados y hábitat asociados en el cantón de la fortuna, San Carlos, Costa Rica. Biocenosis. 2016; 30. Available from: https://revistas.uned.ac.cr/index.php/biocenosis/article/view/1427.

84. Monge Nájera J. Vertebrate mortality on tropical highways: the costa rican case. ida Silv. Neotrop. 1996; 5:154–6. Available from: https://investiga.uned.ac.cr/ecologiaurbana/wp-content/uploads/sites/30/2017/09/JMN-1996-vertebrate_mortality.pdf.

85. Lempp C, Jungwirth N, Grilo ML, Reckendorf A, Ulrich A, van Neer A, et al. Pathological findings in the red fox (Vulpes vulpes), stone marten (Martes foina) and raccoon dog (Nyctereutes procyonoides), with special emphasis on infectious and zoonotic agents in Northern Germany. PLoS One. 2017; 12:e0175469. Epub 2017/04/11. doi: 10.1371/journal.pone.0175469 PMID: 28399176.

86. Navas-Suárez PE, Díaz-Delgado J, Fernandes-Santos RC, Testa-José C, Silva R, Sansone M, et al. Pathological Findings in Lowland Tapirs (Tapirus terrestris) Killed by Motor Vehicle Collision in the Brazilian Cerrado. J Comp Pathol. 2019; 170:34–45. Epub 2019/06/12. doi: 10.1016/j.jcpa.2019.05.004 PMID: 31375157.

87. Navas-Suárez PE, Sacristán C, Díaz-Delgado J, Yogui DR, Alves MH, Fuentes-Castillo D, et al. Pulmonary adiaspiromycosis in armadillos killed by motor vehicle collisions in Brazil. Sci Rep. 2021; 11:272. Epub 2021/01/11. doi: 10.1038/s41598-020-79521-6 PMID: 33432031.

88. Navas-Suárez P. Características e possíveis fatores de risco em cervos neotropicais com histórico de trauma e encaminhados ao Laboratório de Patologia Comparada de Animais Selvagens – LAPCOM, FMVZ, USP, Brasil. Revista de Educação Continuada em Medicina Veterinária e Zootecnia do CRMV-SP. 2016; 14:51–2. Available from: https://www.revistamvez-crmvsp.com.br/index.php/recmvz/article/view/31101.

89. Nettles VF, Quist CF, Lopez RR, Wilmers TJ, Frank P, Roberts W, et al. Morbidity and mortality factors in key deer (Odocoileus virginianus clavium). J Wildl Dis. 2002; 38:685–92. doi: 10.7589/0090-3558-38.4.685 PMID: 12528433.

90. Calero-Bernal R, Pérez-Martín JE, Reina D, Serrano FJ, Frontera E, Fuentes I, et al. Detection of Zoonotic Protozoa Toxoplasma gondii and Sarcocystis suihominis in Wild Boars from Spain. Zoonoses Public Health. 2016; 63:346–50. Epub 2015/11/25. doi: 10.1111/zph.12243 PMID: 26604045.

91. Cantlay JC, Ingram DJ, Meredith AL. A Review of Zoonotic Infection Risks Associated with the Wild Meat Trade in Malaysia. Ecohealth. 2017; 14:361–88. Epub 2017/03/22. doi: 10.1007/s10393-017-1229-x PMID: 28332127.

92. Karesh WB, Noble E. The bushmeat trade: increased opportunities for transmission of zoonotic disease. Mt Sinai J Med. 2009; 76:429–34. doi: 10.1002/msj.20139 PMID: 19787649.

93. Milton AAP, Agarwal RK, Bhuvana Priya G, Saminathan M, Aravind M, Reddy A, et al. Prevalence and molecular typing of Clostridium perfringens in captive wildlife in India. Anaerobe. 2017; 44:55–7. Epub 2017/02/01. doi: 10.1016/j.anaerobe.2017.01.011 PMID: 28159707.

94. Rodriguez-Villar S, Fife A, Baldwin C, Warne RR. Antibiotic-resistant hypervirulent Klebsiella pneumoniae causing community-acquired liver abscess: an emerging disease. Oxf Med Case Reports. 2019; 2019:omz032. Epub 2019/05/31. doi: 10.1093/omcr/omz032 PMID: 31198568.

95. Greig J, Rajić A, Young I, Mascarenhas M, Waddell L, LeJeune J. A scoping review of the role of wildlife in the transmission of bacterial pathogens and antimicrobial resistance to the food Chain. Zoonoses Public Health. 2015; 62:269–84. Epub 2014/08/30. doi: 10.1111/zph.12147 PMID: 25175882.

96. Heaton CJ, Gerbig GR, Sensius LD, Patel V, Smith TC. Staphylococcus aureus Epidemiology in Wildlife: A Systematic Review. Antibiotics (Basel). 2020; 9. Epub 2020/02/18. doi: 10.3390/antibiotics9020089 PMID: 32085586.

97. Mulani MS, Kamble EE, Kumkar SN, Tawre MS, Pardesi KR. Emerging Strategies to Combat ESKAPE Pathogens in the Era of Antimicrobial Resistance: A Review. Front Microbiol. 2019; 10:539. Epub 2019/04/01. doi: 10.3389/fmicb.2019.00539 PMID: 30988669.

98. Tacconelli E, Carrara E, Savoldi A, Harbarth S, Mendelson M, Monnet DL, et al. Discovery, research, and development of new antibiotics: the WHO priority list of antibiotic-resistant bacteria and tuberculosis. The Lancet Infectious Diseases. 2018; 18:318–27. doi: 10.1016/S1473-3099(17)30753-3.

99. Blanco-Peña K, Esperón F, Torres-Mejía AM, La Torre A de, La Cruz E de, Jiménez-Soto M. Antimicrobial Resistance Genes in Pigeons from Public Parks in Costa Rica. Zoonoses Public Health. 2017; 64:e23–e30. Epub 2017/02/24. doi: 10.1111/zph.12340 PMID: 28233464.

100. Hongoh V, Hoen AG, Aenishaenslin C, Waaub J-P, Bélanger D, Michel P. Spatially explicit multi-criteria decision analysis for managing vector-borne diseases. Int J Health Geogr. 2011; 10:70. Epub 2011/12/29. doi: 10.1186/1476-072X-10-70 PMID: 22206355.

101. Grillet ME, Hernández-Villena JV, Llewellyn MS, Paniz-Mondolfi AE, Tami A, Vincenti-Gonzalez MF, et al. Venezuela’s humanitarian crisis, resurgence of vector-borne diseases, and implications for spillover in the region. The Lancet Infectious Diseases. 2019; 19:e149–e161. doi: 10.1016/S1473-3099(18)30757-6.

102. World Health Organization. A global brief on vector-borne diseases. Switzerland: WHO 2014. Available from: https://apps.who.int/iris/bitstream/handle/10665/111008/WHO_DCO_WHD_2014.1_eng.pdf.

103. Dujardin J-C, Herrera S, do Rosario V, Arevalo J, Boelaert M, Carrasco HJ, et al. Research priorities for neglected infectious diseases in Latin America and the Caribbean region. PLoS Negl Trop Dis. 2010; 4:e780. Epub 2010/10/26. doi: 10.1371/journal.pntd.0000780 PMID: 21049009.

104. Montenegro VM, Bonilla MC, Kaminsky D, Romero-Zúñiga JJ, Siebert S, Krämer F. Serological detection of antibodies to Anaplasma spp., Borrelia burgdorferi sensu lato and Ehrlichia canis and of Dirofilaria immitis antigen in dogs from Costa Rica. Vet Parasitol. 2017; 236:97–107. Epub 2017/02/14. doi: 10.1016/j.vetpar.2017.02.009 PMID: 28288773.

105. Rojas A, Rojas D, Montenegro VM, Baneth G. Detection of Dirofilaria immitis and other arthropod-borne filarioids by an HRM real-time qPCR, blood-concentrating techniques and a serological assay in dogs from Costa Rica. Parasit Vectors. 2015; 8:170. Epub 2015/03/23. doi: 10.1186/s13071-015-0783-8 PMID: 25851920.

106. Brenes R, Beaver PC, Monge E, Zamora L. Pulmonary dirofilariasis in a Costa Rican man. Am J Trop Med Hyg. 1985; 34:1142–3. doi: 10.4269/ajtmh.1985.34.1142 PMID: 3834799.

107. Rodríguez B, Arroyo R, Caro L, Orihel TC. Human dirofilariasis in Costa Rica. A report of three new cases of Dirofilaria immitis infection. Parasite. 2002; 9:193–5.

108. Olivo CA, Gundacker ND, Murillo J, Weiss SD, Suarez D, Suarez JA. Subcutaneous Dirofilariasis in a Returning Traveler From Costa Rica. Infect Dis Clin Pract. 2019; 27:58–60. doi: 10.1097/IPC.0000000000000679.

109. Segura NA, Muñoz AL, Losada-Barragán M, Torres O, Rodríguez AK, Rangel H, et al. Minireview: Epidemiological impact of arboviral diseases in Latin American countries, arbovirus-vector interactions and control strategies. Pathog Dis. 2021; 79. doi: 10.1093/femspd/ftab043 PMID: 34410378.

110. Calderón-Arguedas O, Troyo A, Solano ME, Avendaño A, Beier JC. Urban mosquito species (Diptera: Culicidae) of dengue endemic communities in the Greater Puntarenas area, Costa Rica. Rev Biol Trop. 2009; 57:1223–34. doi: 10.15517/rbt.v57i4.5459 PMID: 20073347.

111. León B, Käsbohrer A, Hutter SE, Baldi M, Firth CL, Romero-Zúñiga JJ, et al. National Seroprevalence and Risk Factors for Eastern Equine Encephalitis and Venezuelan Equine Encephalitis in Costa Rica. J Equine Vet Sci. 2020; 92:103140. Epub 2020/06/02. doi: 10.1016/j.jevs.2020.103140 PMID: 32797803.

112. Johnson G, Nemeth N, Hale K, Lindsey N, Panella N, Komar N. Surveillance for West Nile virus in American white pelicans, Montana, USA, 2006-2007. Emerging Infect Dis. 2010; 16:406–11. doi: 10.3201/eid1603.090559 PMID: 20202414.

113. Johnson G, Panella N, Hale K, Komar N. Detection of West Nile virus in stable flies (Diptera: Muscidae) parasitizing juvenile American white pelicans. J Med Entomol. 2010; 47:1205–11. doi: 10.1603/ME10002 PMID: 21175073.

114. Kennedy JM, Earle JAP, Omar S, Abdullah H, Nielsen O, Roelke-Parker ME, et al. Canine and Phocine Distemper Viruses: Global Spread and Genetic Basis of Jumping Species Barriers. Viruses. 2019; 11. Epub 2019/10/14. doi: 10.3390/v11100944 PMID: 31615092.

115. Duque-Valencia J, Sarute N, Olarte-Castillo XA, Ruíz-Sáenz J. Evolution and Interspecies Transmission of Canine Distemper Virus-An Outlook of the Diverse Evolutionary Landscapes of a Multi-Host Virus. Viruses. 2019; 11. Epub 2019/06/26. doi: 10.3390/v11070582 PMID: 31247987.

116. Rendon-Marin S, Martinez-Gutierrez M, Suarez JA, Ruiz-Saenz J. Canine Distemper Virus (CDV) Transit Through the Americas: Need to Assess the Impact of CDV Infection on Species Conservation. Front Microbiol. 2020; 11:810. Epub 2020/05/21. doi: 10.3389/fmicb.2020.00810 PMID: 32508760.

117. Piche M, Alfaro A, Jiménez-Soto M, Murcia P, Jiménez C. Caracterización molecular de dos brotes de distemper canino en animales de vida silvestre en Costa Rica. Ciencias Veterinarias. 2019; 36:38. doi: 10.15359/rcv.36-3.27.

118. Kapil S, Yeary TJ. Canine distemper spillover in domestic dogs from urban wildlife. Vet Clin North Am Small Anim Pract. 2011; 41:1069–86. doi: 10.1016/j.cvsm.2011.08.005 PMID: 22041204.

119. Viana M, Cleaveland S, Matthiopoulos J, Halliday J, Packer C, Craft ME, et al. Dynamics of a morbillivirus at the domestic-wildlife interface: Canine distemper virus in domestic dogs and lions. Proc Natl Acad Sci U S A. 2015; 112:1464–9. Epub 2015/01/20. doi: 10.1073/pnas.1411623112 PMID: 25605919.

120. Beineke A, Baumgärtner W, Wohlsein P. Cross-species transmission of canine distemper virus-an update. One Health. 2015; 1:49–59. Epub 2015/09/13. doi: 10.1016/j.onehlt.2015.09.002 PMID: 28616465.

121. Badilla X, Pérez-Herra V, Quirós L, Morice A, Jiménez E, Sáenz E, et al. Human rabies: a reemerging disease in Costa Rica. Emerging Infect Dis. 2003; 9:721–3. doi: 10.3201/eid0906.020632 PMID: 12781014.

122. Herrera I, Khan SR, Kaleta EF, Müller H, Dolz G, Neumann U. Serological status for Chlamydophila psittaci, Newcastle disease virus, avian polyoma virus, and Pacheco disease virus in scarlet macaws (Ara macao) kept in captivity in Costa Rica. J Vet Med B Infect Dis Vet Public Health. 2001; 48:721–6. doi: 10.1046/j.1439-0450.2001.00485.x PMID: 11846016.

123. Hernandez SM, Peters VE, Weygandt PL, Jimenez C, Villegas P, O’Connor B, et al. Do shade-grown coffee plantations pose a disease risk for wild birds. Ecohealth. 2013; 10:145–58. Epub 2013/05/02. doi: 10.1007/s10393-013-0837-3 PMID: 23636482.

124. Moreno-Díaz M-L, Alfaro E. Valoración socioeconómica del impacto de la variabilidad climática sobre la pesca artesanal en Costa Rica. RU. 2018; 32:18. doi: 10.15359/ru.32-1.2.

125. Choc-M LF, Jiménez-R A, Arguedas-C D, Dolz G. Contracaecum multipapillatum (Nematoda: Anisakidae) en peces de aguas continentales de Guanacaste, Costa Rica e Izabal, Guatemala. Rev Colombiana Cienc Anim RECIA. 2020:e767. doi: 10.24188/recia.v12.n2.2020.767.

126. Mesén Ramírez P, Calvo Fonseca N. Diagnóstico de la Angiostrongilosis abdominal en Costa Rica. Costa Rica: Instituto Costarricense de Investigación y Enseñanza en Nutrición y Salud 2010. Available from: https://www.inciensa.sa.cr/vigilancia_epidemiologica/informes_vigilancia/2010/Parasitologia/Informe%20diagnostico%20Angiostrongiliosis%202010.pdf.

127. Vargas M. Informe técnico Evaluación de test de Morera según resultados del Centro Nacional de Referencia de Parasitología-Inciensa. Costa Rica enero 2012 - abril 2020. Costa Rica: Instituto Costarricense de Investigación y Enseñanza en Nutrición y Salud. 2020. Available from: https://www.inciensa.sa.cr/vigilancia_epidemiologica/informes_vigilancia/2020/CNR%20Parasitologia/Informe%20Tecnico%20A.%20costarricensis.pdf.

128. Santoro M, Alfaro-Alarcón A, Veneziano V, Cerrone A, Latrofa MS, Otranto D, et al. The white-nosed coati (Nasua narica) is a naturally susceptible definitive host for the zoonotic nematode Angiostrongylus costaricensis in Costa Rica. Vet Parasitol. 2016; 228:93–5. Epub 2016/08/26. doi: 10.1016/j.vetpar.2016.08.017 PMID: 27692339.

129. Stunkard. H. W. New intermediate hosts in the life cycle of prosthenorchis elegans (diesing, 1851), an acanthocephalan parasite of primates. J Parasitol. 1965; 51:645–9.

130. Mathison BA, Bishop HS, Sanborn CR, Dos Santos Souza S, Bradbury R. Macracanthorhynchus ingens Infection in an 18-Month-Old Child in Florida: A Case Report and Review of Acanthocephaliasis in Humans. Clin Infect Dis. 2016; 63:1357–9. Epub 2016/08/07. doi: 10.1093/cid/ciw543 PMID: 27501844.

131. Lipkin WI. The changing face of pathogen discovery and surveillance. Nat Rev Microbiol. 2013; 11:133–41. Epub 2013/01/03. doi: 10.1038/nrmicro2949 PMID: 23268232.

132. Gardy JL, Loman NJ. Towards a genomics-informed, real-time, global pathogen surveillance system. Nat Rev Genet. 2018; 19:9–20. Epub 2017/11/13. doi: 10.1038/nrg.2017.88. PMID: 29129921.

133. Mishra N, Fagbo SF, Alagaili AN, Nitido A, Williams SH, Ng J, et al. A viral metagenomic survey identifies known and novel mammalian viruses in bats from Saudi Arabia. PLoS One. 2019; 14:e0214227. Epub 2019/04/10. doi: 10.1371/journal.pone.0214227. PMID: 30969980.

134. Gu W, Deng X, Lee M, Sucu YD, Arevalo S, Stryke D, et al. Rapid pathogen detection by metagenomic next-generation sequencing of infected body fluids. Nat Med. 2021; 27:115–24. Epub 2020/11/09. doi: 10.1038/s41591-020-1105-z PMID: 33169017.

